# Long lifespan is maintained by a unique Heat Shock Factor in reproductive ants

**DOI:** 10.1101/2022.04.07.487479

**Authors:** Karl M. Glastad, Julian Roessler, Janko Gospocic, Shelley L. Berger

**Author notes:** To whom correspondence should be addressed: S.L.B.

## Abstract

Among eusocial insects, reproductive females show longer lifespan than non-reproductive female workers despite high genetic similarity. Using an ant species (*Harpegnathos saltator, Hsal*) featuring inducible worker reproduction and correlated extended lifespan, we find that long-lived reproductive individuals (called “gamergates”) show elevated expression of heat shock response (HSR) genes specifically in the absence of heat stress. This HSR gene elevation is driven in part by gamergate-specific constitutive upregulation of a heat shock transcription factor gene most similar to mammalian HSF2 (called *hsal*HSF2). In sterile workers *hsal*HSF2 is bound to DNA only upon heat stress, but in gamergates *hsal*HSF2 binds to DNA in the absence of heat stress, and correlates with caste-biased gene expression of a subset of heat-inducible genes, thus showing natural bias to gamergates. Remarkably, ectopic expression in *D. melanogaster* of *hsal*HSF2 leads to enhanced survival compared to *hsal*HSF1 following heat stress, as well as extended lifespan. Molecular characterization of these longer-lived flies illustrates multiple parallels between long-lived flies and gamergates, underscoring the centrality of *hsal*HSF2 to extended lifespan in gamergates. Hence, ant caste-specific heat stress resilience and excessive longevity is, remarkably, transferrable to flies via a specialized ant heat shock factor, HSF2. These findings reinforce the critical role of proteostasis to health and to aging, and reveal novel mechanisms underlying facultative lifespan extension.

## Introduction

One of the most fascinating and pressing questions in biology relates to determinative factors of lifespan and potential extension of healthspan. Across Metazoa, several conserved features of molecular aging occur as organisms reach the end of their natural lifespan: changes in the fidelity of genome and epigenome regulation, accumulation of oxidative damage, and accumulation of proteome damage^1,2^. Protein misfolding is a ubiquitous consequence of the everyday function of living organisms, however loss of proteostasis (the health of the proteome) with age has emerged as a major factor that leads to negative consequences of aging, with mis-folded protein accumulation at end-of-life in many species^1,3^. The breadth of this aging phenotype is matched by an equivalently broad distribution and diversity of chaperone proteins that function to re-fold misfolded proteins^1,2^. Further, multiple studies point to up-regulation of chaperone proteins to be a strategy for lifespan extension in mammals^4^, *D. melanogaster*^5,6^ and *C. elegans*^7^.

Early understanding of proteostasis and chaperone function emerged from discovery of the classic heat shock response (HSR)^8,9^, an exceedingly rapid gene induction system. Central to the HSR are the heat shock transcription factors (HSFs), which are DNA binding proteins that induce expression of numerous chaperone proteins in response to proteomic insult. HSF1 is highly studied, and in basal non-heat shock conditions HSF1 is typically sequestered in the cytoplasm as a monomer, bound to the chaperone HSP90 (among others). Upon proteomic insult, HSP90 disassociates from HSF1 monomers, which enter the nucleus, trimerize, and bind to HSR genes, which are rapidly upregulated to combat aggregation of misfolded proteins^10^.

In mammals, multiple distinct HSF transcription factors exist, and play distinct roles outside of this classical mechanism of HSR gene induction utilizing HSF1. For example, in contrast to HSF1 which is ubiquitously expressed, HSF2 shows distinct expression patterns through development^10^, with DNA binding and linked target gene induction showing a more linear correlation with HSF2 expression levels^10^. Furthermore, HSF2 trimerizes with HSF1 monomers, potentially ‘drawing down’ cytoplasmic HSF1 into the nucleus for HSR gene induction independent of strong proteomic insult^11-13^.

Here, we utilize the ant *Harpegnathos saltator* as a non-model long-lived animal, exhibiting inducible lifespan extension. Indeed, while reproductive ants generally have a greatly extended lifespan relative to worker ants, *H. saltator* workers can be naturally or experimentally induced to become reproductive (such reproductively-activated workers are called “gamergates”), and, remarkably, then gain considerable, >6-fold lifespan extension. Because of this inducible lifespan increase, *H. saltator* represents an outstanding model to study aging. Here, we find that, along with lifespan extension, *H. saltator* gamergates possess a much higher resilience to heat stress-induced mortality compared to workers. At the basal level without heat stress, gamergates upregulate a Heat Shock Factor with similarity to HSF2; binding of HSF2 occurs exclusively in gamergates at chaperone genes more highly expressed in gamergates at the basal level without heat stress. Ectopic expression of ant HSF2 in the fly *D. melanogaster* causes resilience to heat stress, extension of fly lifespan, and transcription similar to gamergates without heat stress. Thus, we have linked the remarkable ant caste-specific excessive longevity to the function of a specialized heat shock factor, HSF2, and find that heat resilience and associated long life span can be transferred to flies via expression of HSF2.

## Results

### Gamergates show buffering against heat stress-induced mortality

We tested whether age-matched workers and gamergates can survive prolonged heat stress (18h, 36.5C). Gamergates were considerably more robust to heat stress-induced mortality than workers (∼70% vs ∼40% survival) (Fig 1A). Furthermore, while gamergate survival of heat stress did not differ between younger and older individuals, older workers exhibited a pronounced decrease in survival (∼20%) relative to younger workers (Fig 1A).

**Figure 1.**
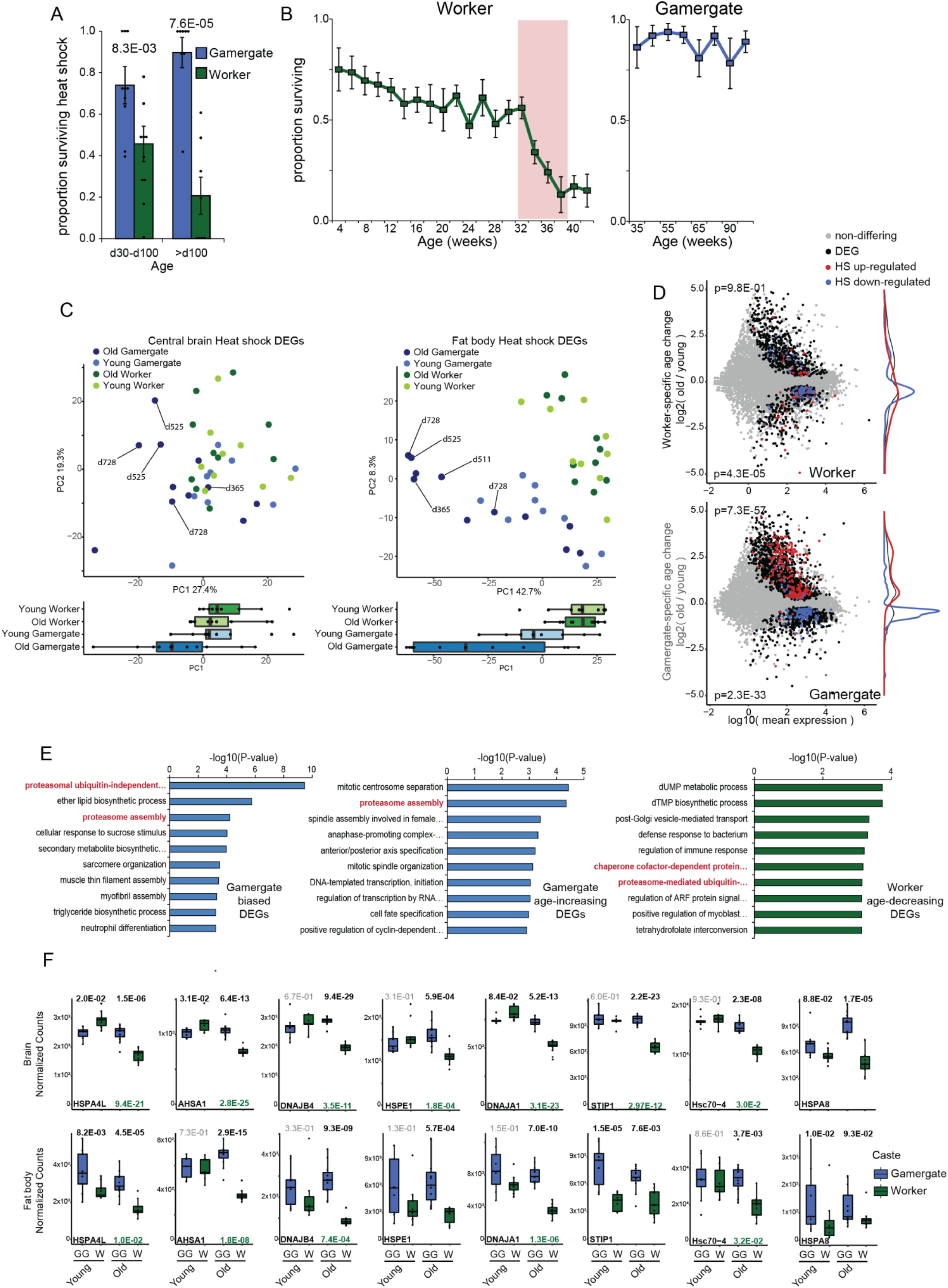
Enhanced heat-shock survival in Gamergates. A) Survival upon prolonged heatshock (18h at 36.5C) for gamergates (blue) and workers (green) of two age groups, showing gamergates show significantly greater survival after heat shock, while workers show age-associated decline in survival. >d100 n=8, d30-d100 n=10. P-values from a Mann-Whitney U test. B) Lifespan of Workers and gamergates. Worker data represent cohorots of ants painted each week and censuseed after one year, then repeated two months later. Gamergate data represent wire-tagged foundational gamergates from the same colonies. C) PCA of aging RNA-seq samples segregated by heat shock up-regulated DEGs comparing data from (left) brains and (right) fat body., illustrating that genes upregulated upon heat shock segregate gamergates and workers (blue and green, respectively), as well as old gamergates from young gamergates (dark and light blue, respectively). D) MA plots of genes showing caste-specific age-related changes in expression, with 1h HS-up and HS-down regulated genes colored in red and blue respectively. P-values in upper and lower are from a fishers exact test comparing up- and down-regulated caste-biased age changing genes to 1h heat shock up- and down-regulated DEGs respectively. E) Gene ontology term (Biological Process) results for the top ten terms enriched among genes (left) biased to Gamergates, (middle) showing significant increase in expression as Gamergates age relative to workers, and (right) showing significant decrease in workers relative to gamergates with age, illustrating enrichment of proteome and chaperone related terms in these categories. F) RNA-seq count plots (normalized counts) illustrating multiple HSPs/HSCs more highly expressed in GGs without heat shock, particularly in older gamergates. P-values represent adjusted P-values from DESeq2.

Because of the known link between proteostasis breakdown and negative consequences of aging ^1^, we assessed whether there is an association between this enhanced survival of gamergates to heat stress (Fig 1A) and previously observed extended lifespan of gamergates relative to workers (Fig 1B)^14^. First, we performed RNA-seq of central brain ^15^ and fat body from workers and gamergates following one hour (40°C) heat stress, focusing on aged ants. We examined responses in in brain because behavior is central within ant systems, and because of the strong link in model organisms between maintenance of proteostasis and neurodegeneration ^16^. Fat body was also utilized due to multiple functions related to aging (liver-like, metabolic functions, adipose tissue) and to its relatively simple structure^17^. We collected age-matched samples of d220-d260 for workers and gamergates, which corresponds to the age of lowest heat stress survival of workers (Fig 1A), and to age-related worker mortality (Fig 1B); we maintained constant age for this comparison between workers and gamergates, despite the considerably longer lifespan of gamergates.

We initially examined the general transcriptomic response to one hour heat stress in both castes together.

Following heat stress we observed 1,425 differentially expressed genes (DEGs) in the fat body (771 upregulated, 654 downregulated; Fig S1A upper panels), and 4.092 DEGs in the brain (2,162 upregulated, 1,930 downregulated; Fig S1B upper panels). The larger number of DEGs detected in the brain is notable, given that neurons in mammals tend to show a blunted HSR^18^. In both tissues, functional terms associated with upregulated genes indicated an immediate HSR to the heat stress insult, including terms related to protein folding and chaperone activity, and to rapid transcriptional response. Downregulated genes indicated more diverse organismal functions, including mRNA processing and translation machinery (Fig S1A and S1B upper panels), as predicted for shut down of normal physiology.

We next assessed caste-specific responses to the one hour heat stress in both tissues to uncover differences that might explain the disparate heat stress survival between castes. We identified DEGs showing significantly stronger up-or downregulation in one caste upon heat stress relative to the other caste (see methods). Surprisingly, despite better survival following the heat stress, there were considerably fewer genes showing gamergate-specific transcriptional response compared to workers (Brain: 803 gamergate-specific DEGs vs 1,188 worker-specific DEGs; Fat body: 91 gamergate-specific DEGs vs 716 worker-specific DEGs; Fig S1A and S1B, lower panels). In fat body, gamergate-specific upregulated DEGs after heat stress showed functional enrichment dominated by chromatin modification and transcriptional regulation (Fig S1A, lower panels), while worker-specific DEGs after heat stress showed top terms similar to general DEGs after heat stress (in both castes); including “protein refolding”, molecular chaperone-related terms, and “response to heat” (Fig S1A, middle panels). In brain, gamergate-specific upregulated DEGs following heat stress showed functional enrichment associated with protein translation as well as HSR (Fig S1B, lower panels), while worker-specific DEGs after heat stress showed stronger enrichment of general HSR pathways (Fig S1B, middle panels). These results indicate that immediately following heat stress, workers, not gamergates, preferentially and more strongly upregulate genes in both tissues related to immediate stress response (including the HSR), which was surprising given the superior survival of gamergates to the heat stress.

### Gamergates naturally up-regulate heat shock response genes and molecular chaperones

Thus in two tissues there were fewer gamergate-specific DEGs related to the HSR compared to worker-specific DEGs at one hour following heat stress despite the considerable survival advantage of gamergates. We hypothesized that gamergates may show a blunted transcriptional response to heat stress because the gamergate transcriptome might exist in a basal state that is more resilient to heat stress. To uncover basal genes specific to gamergates compared to workers, we performed RNA-seq of untreated young (d40-d55) and older ants, in the same two tissues as utilized for the heat stress RNA-seq above. Older workers were aged 220-260 days, which corresponds to the end-of-life of workers (Fig 1B) and matches the ages used for the heat stress analysis above. Older gamergates were aged >280 days.

We first analyzed transcription status within the natural aged ants of the general heat stress upregulated genes (not caste-specific heat stress upregulated) analyzed above. We performed PCA analysis on the natural aging samples, to assess whether genes upregulated upon heat stress exhibited differential expression between gamergates and workers during aging. In fat body, heat stress upregulated genes strongly segregated the transcriptome of natural aged workers and gamergates (Fig 1C, right), showing that, heat stress DEGs vary in expression in a caste-specific manner. Specifically, while this pattern of genes responding to heat stress segregating aging caste was less pronounced in brain (Fig 1C, left), strikingly, in both tissues older gamergates very strongly segregated on the PCA while older workers did not (Fig 1C). These results indicate that both gamergate status and gamergate age are associated with genes differentially regulated in response to heat stress.

We then directly compared age-associated DEGs of natural castes to DEGs upon heat stress. Overlapping heat stress DEGs with gamergate and worker-specific age DEGs in fat body revealed that genes that change preferentially in gamergates with age showed striking correspondence to genes differentially regulated upon heat stress (Fig 1D, lower). This striking relationship either did not appear at all, as seen among fat body worker-specific age DEGs (heat stress upregulated DEGs), or existed to a much weaker degree (heat stress downregulated DEGs; Fig 1D, upper). In the brain, we also observed a consistent relationship between gamergate downregulated genes and heat stress downregulated genes, but not for upregulated genes (Fig S1C, lower). Notably, workers displayed an opposite trend for heat stress upregulated genes: heat stress upregulated genes significantly overlapped with genes decreasing specifically in workers with age (Fig S1C, upper). This latter finding suggests that expression of HSR genes may decrease in worker brains as they age.

In fat body, we observed enrichment of functional terms related to the proteasome or protein chaperones among genes generally biased to gamergates (Fig 1E, left), among genes preferentially increasing in gamergates with age (Fig 1E, middle), and among genes preferentially decreasing in workers with age (Fig 1E, right). These gene expression patterns indicate that regulation of the proteome and proteostasis distinguishes the differential caste and aging transcriptome programs of gamergates and workers. In brain, despite the lack of global association between heat stress upregulation and gamergate-specific age-increasing genes, we observed a similar pattern among genes biased to old gamergates, which showed strong enrichment for chaperone and protein folding terms (Fig S1D, lower). Consistent with our prior result, genes decreasing in worker brains with age were strongly enriched for similar terms (Fig S1D, upper). In summary, in both brain and fat body, aging gamergates upregulate a HSR gene expression program, although in brain this is less clear on the global level. Importantly, in both tissues, workers downregulate heat shock and protein folding genes with age, while gamergates do not (Fig 1D and E; Fig S1C and D). These gene expression observations are strongly consistent with the stress resilience and long lifespan of gamergates relative to workers.

One of the main components of heat stress-related mortality is aggregation of misfolded proteins^3,19^, and chaperone proteins help to correct this^20-22^. We thus examined the natural aging DEGs for annotation of heat shock or chaperone proteins. In both tissues, we identified numerous genes biased to gamergates that manage mis-folded proteins (Fig 1E and Fig S1D), consistent with our functional enrichment results. Indeed, of well-known chaperone genes conserved between ants and mammals, we found 18 genes that were significantly higher in at least one tissue in gamergates (Fig S2 A and B); 17 genes showed significant bias to old gamergates in fat body, and 13 in brain (Fig S2A and B). Ten of these were highly consistent between brain and fat body (top 8 shown in Fig 1F). Interestingly, in brain, worker expression of these genes decreased with age (Fig 1F and S2A). These genes included HSP, DNAJ, and alpha-crystallin proteins, which have central roles as molecular chaperones to manage protein folding and misfolding ^23,24^. In summary, aging gamergates upregulate key gene expression pathways to maintain protein homeostasis, while workers downregulate these pathways.

### HSF2 shows gamergate-biased expression without heat shock

We sought to understand whether upstream transcriptional regulators might drive this remarkable pattern of gamergate-biased stress gene expression in the basal state. Heat shock factors (HSFs), a set of DNA-binding transcription factors, famously upregulate HSR genes encoding heat shock proteins (HSPs) including molecular chaperones in classic heat shock pathways and in response to other stressors. We uncovered an HSF transcription factor paralogue in *H. saltator* (as well as in multiple other Hymenoptera) that does not exist in *D. melanogaster*. Strikingly, this HSF gene is markedly upregulated in the fat body of old gamergates while decreasing in expression in workers (Fig 2A, lower); in brain, this HSF is slightly elevated in old gamergates but markedly lowered in expression in old workers (Fig 2A, upper). We aligned this protein, and the annotated *H. saltator* caste-invariant HSF1 (Fig 2A, left), to *H. sapiens* HSF proteins, which revealed similarity to human HSF2 protein (Fig 2B). In mammals, HSF2 plays a heat shock-independent role during development, but also upregulates HSR genes similarly to HSF1 ^11^; this suggests that *H. saltator* HSF2 may play a secondary role of activating stress-response genes at the basal state in gamergates, independent of the classical HSR pathway of HSF1. Curiously, HSF2 is a truncated form of typical HSFs, lacking the TAD (transcriptional activation domain) and RD (regulatory domain), but notably maintaining an intact DNA binding domain. HSF2 also possesses a partial oligemerization domain (HR-A/B) (Fig 2B, bottom), however given the low conservation of HR-A/B, it is unclear if HSF2 is able to trimerize with HSFs. HSF2 was present in the genome of all hymenopteran species examined, illustrating its evolutionary conservation across eusocial and non-eusocial species (Fig S2C).

**Figure 2.**
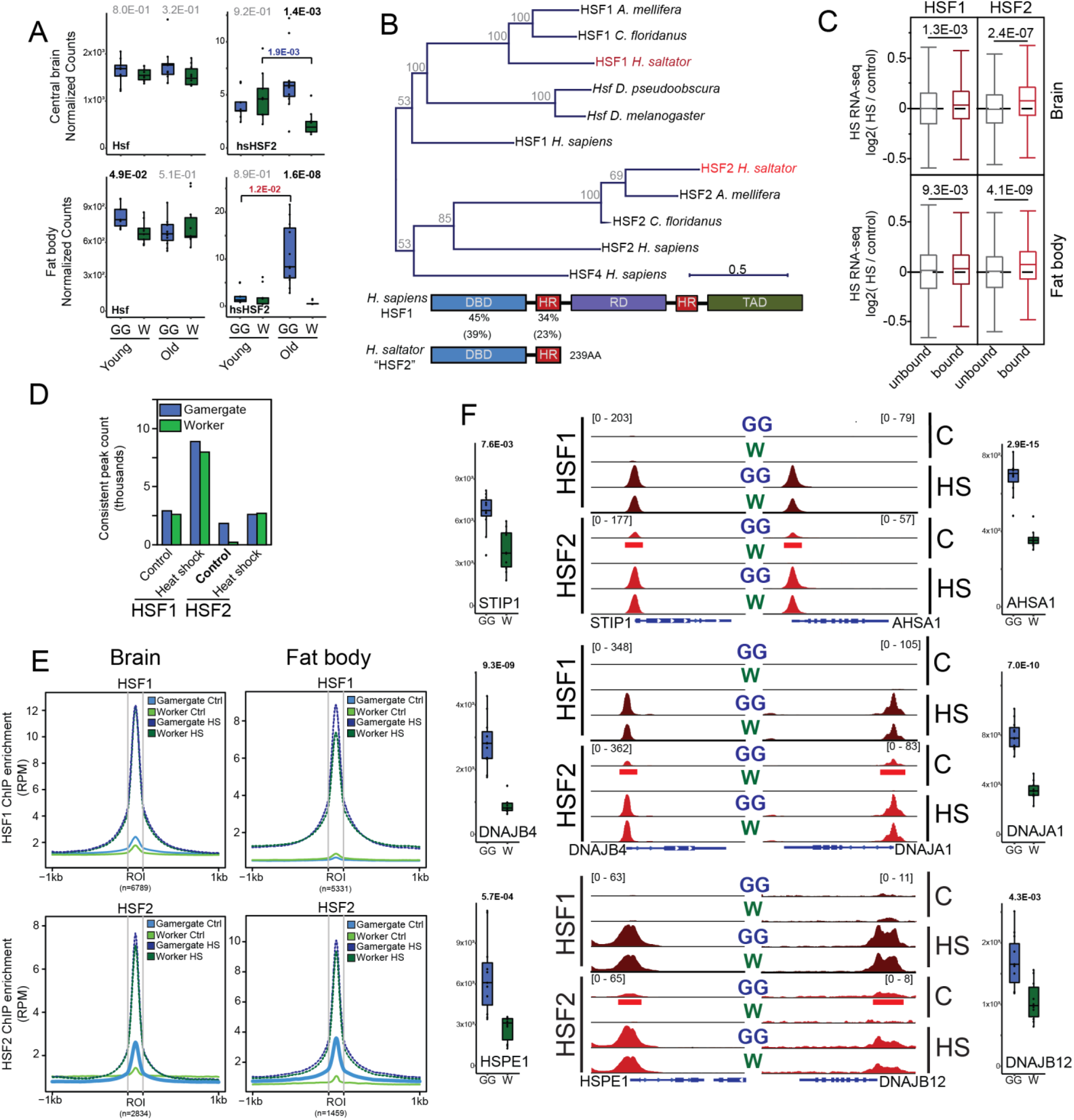
HSF2 shows gamergate-biased expression and non-heat shock binding. A) Normalized counts for HSF1 (left) and HSF2 (right) in *H. saltator* young and old workers and gamergates for (top) brain and (bottom) fat body, showing gamergate-biased expression of HSF2 in older ants. P-values represent adjusted p-values taken from DESeq2. For HSF2 genes showing (blue p-values) age related decline or (red) age related increase in expression are given when significant. B) Protein alignment (PRANK on protein sequences) phylogenetic tree illustrating that HSF2 is closer to *H, sapiens* HSF2 relative to HSF1 and HSF4. Numbers given at nodes represent bootstrap values after 1000 iterations. Below is given a schematic of *H. sapiens* HSF1 as compared to *H. saltator* HSF2, with percent AA conservation given between respective domains (parenthetical values represent percent conservation as compared to *D. melanogaster Hsf*). C) HSF2 (light red) is more predictive of heat shock up-regulation of gene expression than HSF1 (dark red) for both 1h post-heat shock (top) and 18h post heat-shock (bottom). P-values from a mann-whitney U test comparing genes lacking HSF binding to those possessing it. D) Number of peaks detected in data from fat body for HSF1 and HSF2 in worker and gamergate, illustrating very low peak count for HSF2 in worker without heat shock (Control), while non-heat shocked gamergates show similar numbers of peaks with and without heat shock. E) Meta plots of normalized ChIP signal (RPM) for HSFs at peaks generally bound by the respective HSF upon heat shock, showing that HSF2 (right) shows binding in gamergates only (light blue) without heat shock at loci bound by HSF2 upon heat shock (irrespective of caste). F) Example genome browser tracks for 6 gnees shown in Fig 1F illustrating binding by HSF2 in gamergates in the absence of heat shock (red underlined peaks). For respective HSFs, all tracks were autoscaled to one another (RPM values given in brackets). C: Control, HS: heat shock, GG: gamergate samples, W: worker samples. Aside each track is given the count plots representing old caste untreated RNA-seq (as shown in Fig 1F). P-values represent adjusted p-values from DESeq2.

To test whether HSF2 specifically activates stress-response genes in young gamergates, we developed antibodies to *H. saltator* HSF1 and HSF2 (Fig S2D). There were constant protein levels of HSF1 and elevation of HSF2 in gamergates relative to workers (Fig S2D) similar to RNA levels (Fig 2A). We performed ChIP-seq of HSF1 and HSF2 in gamergates and workers with heat stress and without stress, focusing on the same ages as for heat stress RNA-seq above. We found strong binding of both HSF1 and HSF2 in fat body and brain with heat stress (Fig S2E and S2G), as expected given their known roles in mediating the HSR, with >7,000 sites bound consistently among replicates by HSF1 upon heat stress and 2,612 sites by HSF2 (Fig S2E). The majority of sites bound by HSF2 upon heat stress were also bound by HSF1, with HSF1 targeting more genes than HSF2 (Fig S2E). Genes bound by HSF1 and HSF2 during heat stress were generally characterized by functional terms related to translation, protein folding, and heat response. Importantly, in gamergates, we found far more sites bound by HSF2 without heat stress, (Fig S2E), which may reflect a non-standard role in model system neurons^18^. Hence, HSF1 functions similarly in gamergate and worker in heat stress conditions, whereas in non-heat stress basal conditions, HSF2 shows strong binding within the gamergate genome, but not within the worker genome.

We then examined the overall role of HSF1 and HSF2 in binding during the heat stress transcriptional response. Surprisingly, when comparing HSF ChIP-seq binding to heat stress RNA-seq, we found that HSF2 binding was more predictive of general heat stress upregulation of gene expression, for both brain (Fig 2C upper) and fat body (Fig 2C lower) RNA-seq, illustrated in tracks taken from worker (FigS2G). We compared HSF peak enrichment to log2 fold difference of RNA-seq between heat stress and control for both tissues, and found HSF2 enrichment was also more strongly correlated with level of heat stress gene induction (although HSF1 was also predictive of heat stress gene upregulation; Fig S2F). Because HSF1 binds the majority of loci bound by HSF2, this stronger correspondence between HSF2 and heat stress gene expression upregulation may be due to HSF2 showing more specificity to direct the upregulation of heat stress genes. In support of this, correlations between heat stress gene expression change and HSF1 peak enrichment were stronger when only considering genes also marked with HSF2 (Fig S2F, right). This suggests that HSF1 is more permissive in its targeting, including either genes showing weaker heat stress response, or including genes that are repressed upon heat stress, as seen in other systems ^25^.

### HSF2 binds genes in gamergates without heat stress

As mentioned above, we observed a striking difference in HSF2 ChIP-seq peak counts for non-heat stress basal condition in gamergate compared to worker (Fig 2D, S2E). Notably, HSF2 showed marked binding in gamergates without heat stress in both tissues (Fig 2E, lower, thick blue line), but much less or no binding in workers without heat stress (Fig 2E lower, thin green line). Thus, HSF2 binding without heat stress in gamergates may poise upregulation of heat stress response genes in aging gamergates, and this may be central to long lifespan in gamergates. To assess this, we examined the cohort of naturally gamergate-biased HSR genes/chaperone protein genes (Fig 1F) for binding of HSF2 without heat stress. We found that 14/18 of these genes showed gamergate-specific binding in the absence of heat stress (six are shown in Fig 2F, and six in Fig S3A; brain: Fig S3B), and these same genes displayed robust binding of both HSF1 and HSF2 in both castes upon heat stress (Figs 2F and S3A, S3B). Indeed, the majority of DNA sites bound by HSF2 in unstressed gamergates became much more highly bound in both castes with heat stress (Fig 2E), implying that HSF2 targeting to genes without heat stress operates under the same principles as with heat stress, but, notably, binding without heat stress is specific to HSF2 binding within gamergates.

Globally, genes showing caste-specific binding of HSF2 in basal non-heat stress conditions, which were all gamergate-biased in HSF2 binding, also showed a pattern of moderate expression bias to gamergates (fat body shown in Fig 3A top). This moderate bias is likely because HSF2 is not the major driver of caste-specfic differential gene expression^26^; instead, we propose that HSF2 mediates gamergate-biased gene expression in the basal condition for a subset of stress response genes, and this is supported by the observation that the magnitude of HSF2 association is considerably higher when limited to genes showing heat stress induction in our data (fat body: Fig S3C; brain Fig S3D). Indeed, among the 216 genes showing general gamergate bias also featuring a non-heat stress HSF2 peak, the functional categories most enriched were associated with HSR or unfolded protein binding.

**Figure 3.**
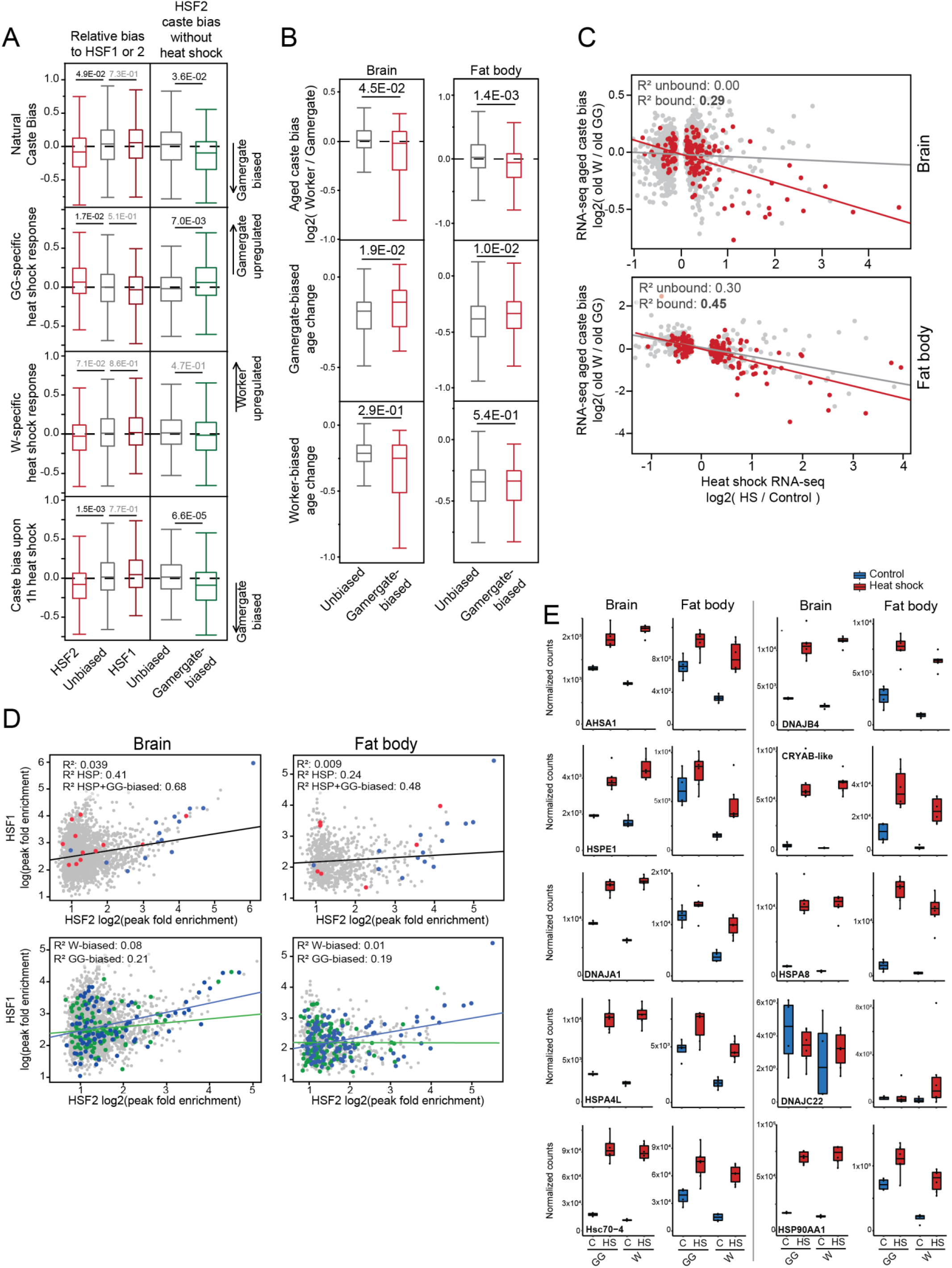
HSF2 in gamergates without heat shock is associated with gamergate-bias in old castes. A) For fat body, heat shock DEGs showing higher relative levels of HSF2 (as compared to HSF1; left plots) as well as those bound by HSF2 in gamergates without heat shock (right plots) show (from top to bottom) significant bias to gamergates, gamergate-biased up-regulation upon heat shock but not worker-biased, as well as gamergate bias among heat shocked samples. P-values from a mann-whitney U test comparing genes without HSF2 or with no difference between HSF2 and HSF1 to the respective categories. B) Genes bound by HSF2 in gamergates without heat shock and also show a consistent pattern of decreased expression with age irrespective of caste, show consistent bias to gamergates (top) and higher expression in aged gamergates but lower expression in aged workers for both brain (left) and fat body data (right). P-values represent results from a mann-whitney U test comparing gamergate-biased HSF2 without HS to unbiased or genes with HSF2 only *with* heat shock. C) Correlation scatterplots comparing general heat shock RNA-seq HS bias to aged caste RNA-seq, divided into genes with (red) and without (grey) HSF2 binding in gamergates without heat shock, illustrating that in both tissues (top: brain; bottom: fat body), HSF2 binding in gamergates is assocated with genes that much more strongly show an association between gamergate-bias and heat shock upregulation. D) correlation scatterplots comparing gene-linked peak enrichment for control (x-axis) HSF2 and (y-axis) control HSF1 in brain (left) and fat body (right), subset to genes with known chaperone function (top row; from Fig 1F and S2) that are (blue) and are not (red) gamergate biased, as well as (bottom) genes showing bias to workers (green) and gamergates (blue), illustrating that among core HSPs as well as genes biased to old gamergates, HSF1 and HSF2 levels show a positive correlation, but outside of these gene categories there is little relationship. E) Known heat-shock related genes from Fig 1F, illustrating that many show larger magnitude of up-regulation in workers, despite many showing as high or higher overall expression in gamergates following 1h of heat shock (light red) in both tissues.

We then examined whether HSF2 binding in gamergates without heat stress correlated with increased expression in old gamergates. We found that in both tissues (upon subsetting our data to genes showing general decline in our aging data irrespective of caste), HSF2 gamergate-bound genes (1) were more biased to old gamergates (Fig 3B, upper), (2) showed increased expression (relative to unbound genes) as gamergates aged (Fig 3B, middle), (3) but showed decreased or non-significant expression as workers age (Fig 3B, lower). This suggests that HSF2 may be acting more strongly in aging gamergates to upregulate or maintain levels of chaperone expression relative to aging workers.

We examined whether HSF2-bound genes are more predictive of the association described above between heat stress gene expression and gamergate bias. Indeed, at heat stress DEGs, genes bound by HSF2 in gamergates without heat stress showed a much stronger association between heat shock differential regulation and caste bias among old gamergates (Fig 3C). Specifically, genes bound by HSF2 in gamergates without heat stress are more likely to be upregulated with heat stress *and* to show bias to old gamergates without heat stress. This further supports our hypothesis that the observed association between gamergate bias and heat stress differential expression is driven, in part, by HSF2 binding in gamergates without heat stess.

We further note that, for many core chaperone genes (e.g. Fig 1F) that show bias to old gamergates, HSF2 expression levels (Fig 2A) track chaperone levels. That is, in fat body, HSF2 is more highly expressed in old gamergates (relative to all other samples) and chaperone genes show a similar pattern of increase in old gamergates, while, in brain, HSF2 decreases in expression for old workers and chaperone gene expression levels decrease in old workers (Fig S2A and B).

Because *H. saltator* HSF2 lacks several functional domains found in mammalian HSF2, we examined whether HSF2 may act, in part, by co-binding with HSF1 in the absence of heat stress, as observed in mammals ^13^. While the overall correlation between HSF1 and HSF2 binding enrichment levels was weak without heat stress, the co-binding was stronger among chaperone genes, with the strongest relationship for chaperone genes that show gamergate bias (Fig 3D, top). Further, genes generally biased to gamergate expression (independent of chaperone function) showed a stronger correlation between basal levels of HSF2 and HSF1 binding than worker-biased genes (Fig 3D, bottom). This suggests that among heat stress responsive genes, HSF2 may mediate increased HSF1 binding, and that this relationship may also exist among a subset of gamergate-biased genes.

Overall, these data are consistent with HSF2 serving to bias expression of a subset of stress response genes to gamergates even in the absence of heat stress, which may in turn explain the stronger differential transcriptional response to heat stress among workers compared to gamergates. Hence, returning to the 18 gamergate-biased core chaperone genes, we found that 8/14 of these showed significantly higher worker-upregulation in fat body with heat stress (as compared to gamergates) (Fig 3E, S2B); indeed, inspection of expression level of these genes upon heat stress revealed that this was not necessarily due to higher overall worker expression with heat stress, but simply that the magnitude of upregulation in workers was higher, and this was due, in almost every gene, to elevated basal expression in unstressed gamergates (Fig 3E).

### Ectopic expression of HSFs extend fly survival under heat shock and stressed aging

To explore the mechanism by which HSF2 buffers *H. saltator* gamergates against stress and extends lifespan, we expressed ant HSF1 and HSF2 (hereafter *hsal*HSF1/2) in the fly *Drosophila melanogaster*. We produced fly lines with UAS-driven ectopic expression of *hsal*HSF1 and *hsal*HSF2. As an HSF overexpression control we utilized flies produced in the same way as our *H. saltator* HSF fly lines, but with similarly expressed *D. melanogaster Hsf* gene (hereafter, *dm*HSF), and additionally, with nuclear RFP as a general overexpression control (Fig S4A for validation). We crossed these flies to elav-Gal4 and PPL-Gal4 driver lines to produce flies expressing these genes in neurons and fat body cells, respectively, to match our ant data.

We evaluated survival under heat stress of varying intensity: 45m at 41C, 4h at 37C and 24h at 35.5C. For most conditions, strikingly, *hsal*HSF2 ectopic expression led to increased survival under heat stress conditions compared to *hsal*HSF1 or to *nRFP* (Fig 4A and 4B). We also noted that, despite much higher similarity of *hsal*HSF1 to fly *dm*HSF, *hsal*HSF1 showed less buffering against heat stress compared to *hsal*HSF2. When expressed in neurons *hsal*HSF1 did not significantly differ from nRFP ectopic expression controls, except in the context of 24h heat stress (Fig 4A, right), while, in fat body, all three HSF ectopic lines showed increased survival relative to the nRFP control; Fig 4B).

**Figure 4.**
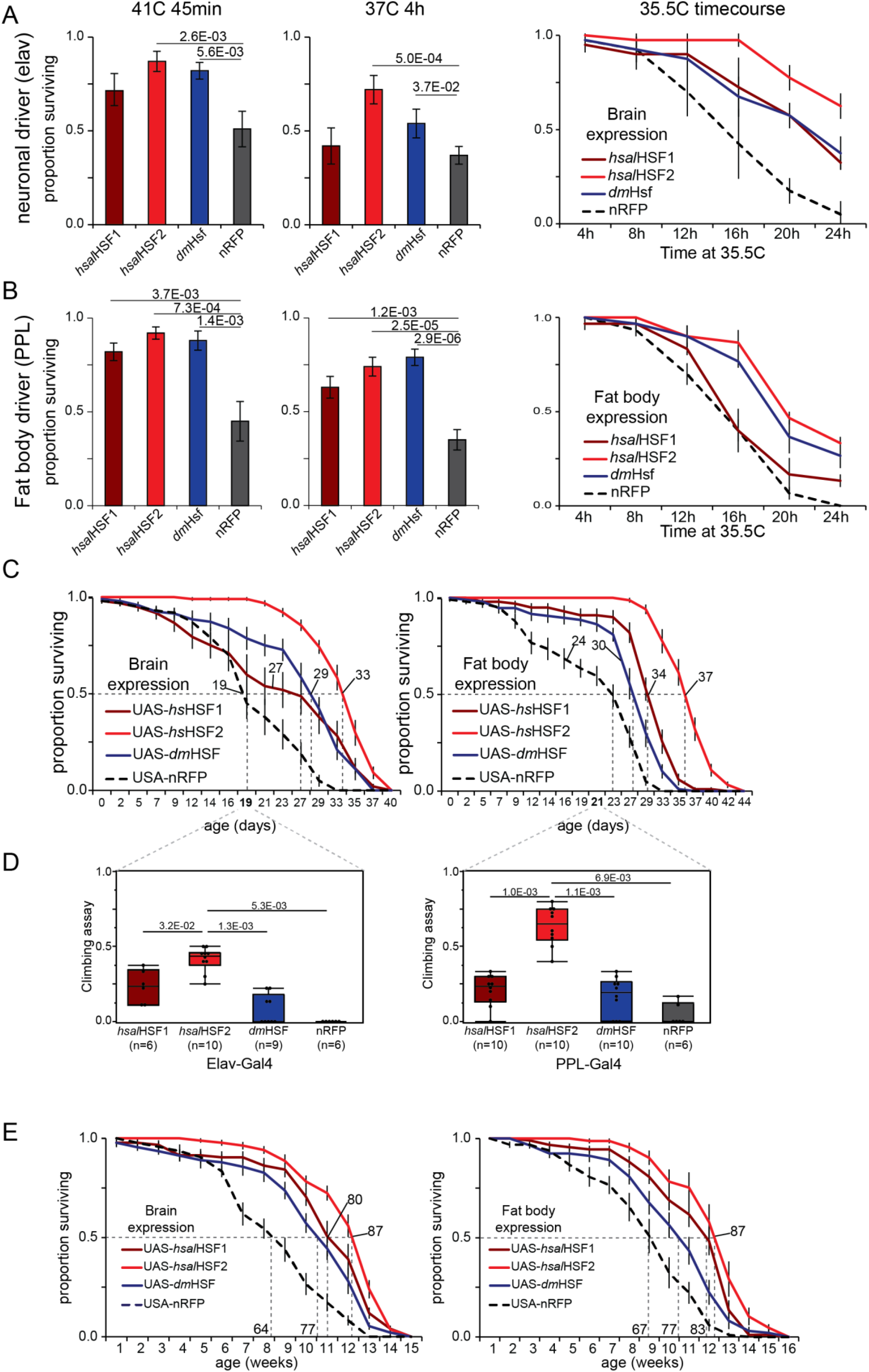
Ectopic expression of HSFs increase survival with heat shock and extend fly lifespan. A) Proportion of flies ectopically expressing *H. saltator* HSF1, *H. saltator* HSF2, *D. melanogaster Hsf*, or identically-transformed nRFP in neurons that survived short-term high heat (41C for 45min; leftmost plot, n=10), medium heat stress (37C for 6h; middle plot, n=10), and 4h timepoints across a survival timecourse under mild heat stress (35.5C; rightmost plot, n=3-4 per timepoint). B) The same as (A) but for fat body driven ectopic expression of the same genes. C) Lifespan assay of flies ectopically expressing *H. saltator* HSF genes, nGFP (Iso31 background), or genetic background control (attp) when kept at 30C for neuronal ectopic expression (left) or fat body ectopic expression (right). Numbers labeling lines represent median lifespans for given curves. D) Results of climbing assay taken from indicated day of 30C aging assay. P-values represent results from a Mann-Whitney U test. E) Results from lifespan assay performed at 25C for ectopic HSF expression lines (or nRFP negative control) driven under (left) elav or (right) PPL drivers. Numbers labeling lines represent median lifespan for given curve.

We performed a short-term stress-associated “aging” assay at 30C, simulating longer-term low-stress conditions. As with the stronger heat stress (Fig 4A, 4B), ectopic expression of HSFs led to longer lifespan, with *dm*HSF and *hsal*HSF1 showing intermediate lifespan extension (median lifespan in days *dm*Hsf elav: 29, PPL: 30; *hsal*HSF1 elav: 27, PPL: 34; Fig 4C). Strikingly, again, *hsal*HSF2 overexpression led to improved survival compared to other ectopic expression lines, showing >50% higher median lifespan in this context relative to nRFP controls (*hsalt*HSF2 median elav: 36 days, PPL: 38 days; Fig 4C).

Additionally, we observed increased locomoter activity in our *hsal*HSF2 ectopic expression flies and performed a common assay to assess fly fitness (climbing assay, see methods) at d19 and d21 (for elav and PPL-driven expression, respectively). The *hsal*HSF2 expressing flies outperformed all other lines in climbing after ∼20 days under mild heat stress (Fig 4D). This was particularly striking for PPL-driven fat body expression, despite the non-neuronal expression of *hsal*HSF2 (Fig 4D, right).

Importantly, in a conventional aging assay, we observed that flies expressing *hsal*HSF2 showed a 30% extension of lifesan (*hsal*HSF2 lifespan: 87 days, both drivers; Fig 4E). Thus, remarkably, compared to *hsal*HSF1 or *dm*HSF, *hsal*HSF2 extension of lifespan operates beyond buffering against heat stress (Fig. 4C), and also plays a role in extending lifespan (Fig 4E), consistent with our observations in *H. saltator*.

### Conserved targets mediate stress-buffering and lifespan extension of hsHSF2 in ant fly

We explored the the mechanism whereby *hsal*HSF2 expression in flies increases survival under stressed and aging non-stressed conditions. We performed RNA-seq of the four ectopic fly lines (*hsal*HSF1, *hsal*HSF2, *dm*Hsf, and nRFP) driven in fat body (PPL) with and without heat shock. Consistent with the distinct survival of *hsal*HSF2 flies, this group showed a distinct transcriptional signature both in the control without heat stress, and after heat stress (Fig 5A). Here we compared each distinct fly line to all others for both control and heat stress conditions. We found that the number of genes significantly higher or lower in *hsal*HSF2 flies compared to all others in a combined framework was remarkably higher than for any other fly line (Fig 5A). Furthermore, examination of functional enrichment of genes more highly expressed in *hsal*HSF2 flies in control without heat stress revealed strong enrichment of functional terms related to protein folding, chaperone activity, and general HSR (Fig 5A, top right red font). Genes preferentially more highly expressed in heat stressed *hsal*HSF2 flies showed enrichment for mitochondria-related functions, chromatin modification, and DNA repair terms (Fig 5A, bottom right)

**Figure 5.**
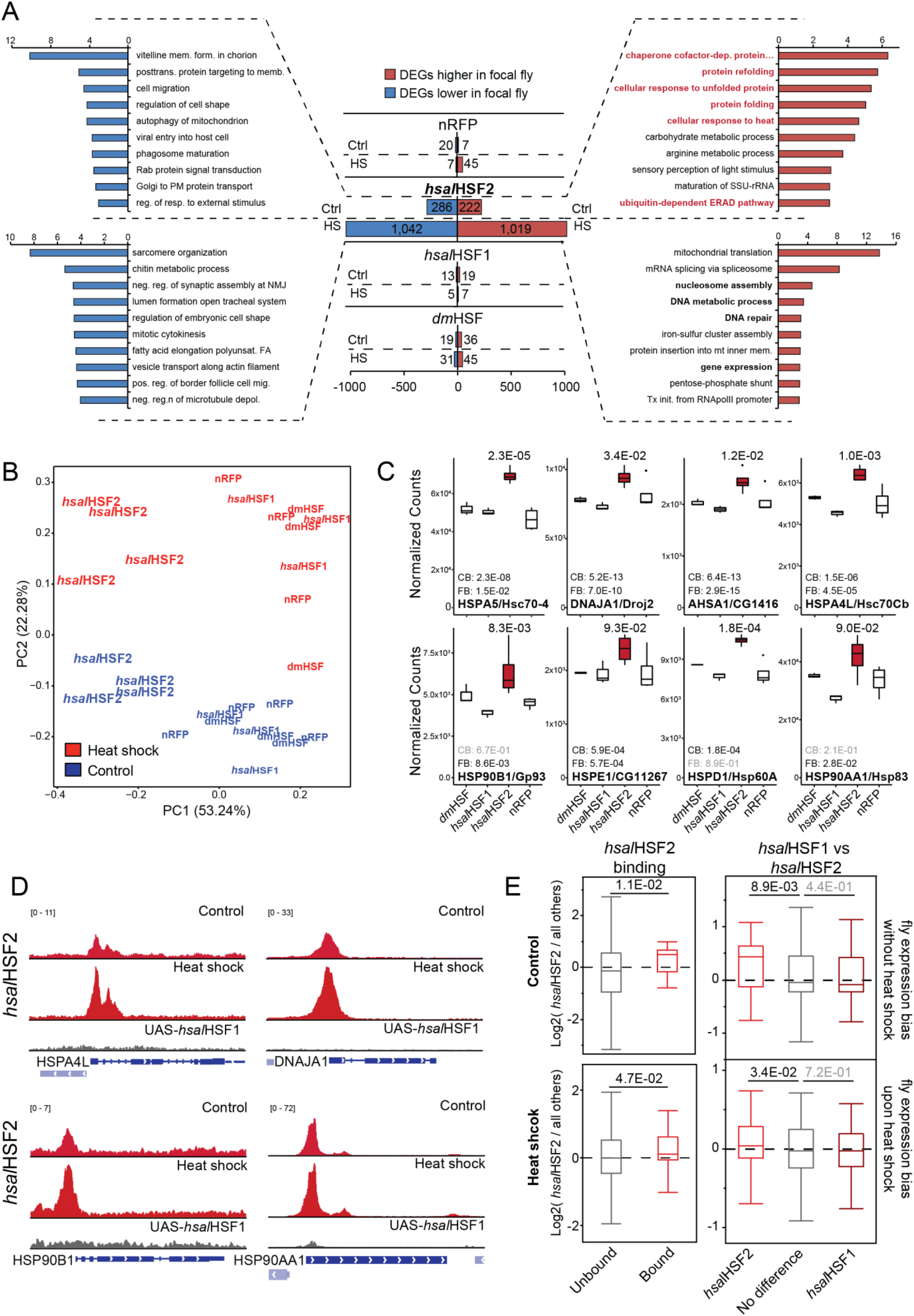
Genome-wide characterization of HSF ectopic expression flies reveals conserved targets and binding with ants. A) Numbers of differentially expressed genes between the given ectopic fly line and all others here for (top bars of each pair) heat shocked samples and (bottom bars for each pair) non-heat shocked samples, illustrating strong departure of HSF2 flies from others shown. Offset bar plots represent top 10 significant gene ontology terms (Biological Process) enriched among the respective groups of HSF2 up-or down-regulated genes. Red terms illustrate the strong presence of genes with functions related to managing mis-folded proteins among HSF2 flies in the *absence* of heat shock, similar to what is seen in natural gamergates. B) Principle component analysis of RNA-seq from all transgenic fly samples illustrating transcriptionally-distinct nature of HSF2 overexpression flies relative to all others. PCA generated using VST-transformed counts of all genes differing significantly (padj < 0.01) in a combined model modeling genetic line and heat shock status. C) Normalized count plots of non-heat shocked fly samples illustrating key chaperone genes most highly expressed in *hsal*HSF2 flies without heat shock that also feature significant gamergate bias in *H. saltator* aging RNA-seq. Values given above gene names represent adjusted p-values taken from *H. saltator* caste comparisons between old gamergates and workers for gamergate-biased genes. CB: central brain adjusted p-values; FB: fat body adjusted p-values. D) Example genome browser tracks from four genes featuring *hsal*HSF2 heightened expression without heat shock as well as *hsal*HSF2 binding in the same context. Given below heat shock and control tracks are tracks from ChIP-seq using the antibody targeting *hsal*HSF2 in heat shocked *hsal*HSF1 flies. E) Boxplots of average log2 expression ratios for DEGs between HSF2 flies and all other fly lines analyzed here, illustrating that (left) HSF2 binding (as determined by peak presence in ChIP-seq of transgenic flies) is predictive of HSF2 fly transcriptional bias, as is (right) higher relative enrichment of HSF2 (vs hsHSF1), both for flies in the absence of heat shock (top) as well as heat shocked samples (bottom). P-values from a Mann-whitney U test comparing genes without HSF2 or with no difference between HSF2 and HSF1 to the respective categories.

This transcriptional difference between *hsal*HSF2 flies and the other flies was underscored by principle component analysis of our fly RNA-seq samples (see methods) showing strong differentiation of *hsal*HSF2 flies relative to all others, regardless of heat stress (Fig 5B). Surprisingly, despite the observation that *dm*HSF and, in some contexts, *hsal*HSF1 ectopic flies showed moderately enhanced survival (Fig 4B and C), in stark contrast to *hsal*HSF2, they did not differ in transcription substantially from one another nor from nRFP flies with or without heat stress (Figs 5A and 5B).

Given this and our results from *H. saltator*, we examined individual genes contributing to the enrichment of chaperone-related functions among basal non-heat stressed *hsal*HSF2 flies. We found that, of the 222 genes more highly expressed among non-heat stressed *hsal*HSF2 flies, 17 were directly annotated with the term “protein folding” and 30 with “response to stress”. We then compared across species to discover key conserved genes activated by ant HSF2 that might regulate heat resilience and lifespan. Over half of the genes annotated with “protein folding” (9/17) were also significantly more highly exressed in at least one tissue in gamergates, and/or upregulated preferentially as gamergates age (7 in FB, 6 in brain; Fig 5C and Fig S4C).

We sought to assess whether ectopic *hsal*HSF2 expression led to this remarkably strong distinct transcription via binding of target genes without heat stress. We performed ChIP-seq of *hsal*HSF1 and *hsal*HSF2 in our fat body (PPL) ectopic expression flies, utilizing the custom antibodies. Of the 17 genes more highly expressed in control *hsal*HSF2 flies vs others, we found that 11 featured an *hsal*HSF2 peak in non-heat shocked samples (four shown in Fig 5D), similar to our ant results. Of the nine showing *hsal*HSF2 fly expression bias and also gamergate bias, eight featured an *hsal*HSF2 peak in fly and all featured an *hsal*HSF2 peak in gamergates without heat shock, implying that many are directly up-regulated by *hsal*HSF2 binding, and, importantly, this mechanism similarly occurs in gamergates.

We noted that, for both *hsal*HSF1 and *hsal*HSF2, binding was observed with and without heat stress, with *hsal*HSF1 showing many more bound loci than *hsal*HSF2 (Fig S5A, S5B), similar to our obervations in *H. saltator* (Fig S3B). Also similar to *H. saltator*, despite the abundance of *hsal*HSF1 binding in our fly data, *hsal*HSF1 binding was less predictive of heat stress upregulation than was *hsal*HSF2 binding (Fig S5C). *hsal*HSF1 also showed lower increase in binding upon heat stress relative to *hsal*HSF2 (Fig S5D). For *hsal*HSF2 this may reflect natural propensity to bind in an expression-associated manner to activate genes as in *H. saltator*. In contrast, however, for *hsal*HSF1 this may be due either to ectopic expression leading to excess *hsal*HSF1, or to protein divergence between *hsal*HSF1 and *dmelHsf* leading to inability of hsp90 to bind to *hsal*HSF1 in non-stressed conditions to retain *hsal*HSF1 in the cytoplasm, as typical for HSF1/*Hsf*. This is supported by the fact that in *H. saltator hsal*HSF1 binds far fewer genes in fat body without heat stress then with heat stress (Fig S2E, right plots). We used an important control for the *hsal*HSF2 fly ChIP-seq, which included a ChIP of *hsal*HSF2 in *hsal*HSF1 flies to ensure the specificity of our *hsal*HSF2 antibody, finding no strong cross-reactivity with *hsal*HSF1 (or endogenous *dmel*HSF; Fig S5B and 5D, grey tracks).

Finally, we found that *hsal*HSF2 binding was also more broadly predictive of expression bias in fly (Fig 5E, left; Fig S5E, left bottom), consistent with the hypothesis that *hsal*HSF2 fly lifespan and stress resistence was partially due to specific *hsal*HSF2 binding. *hsal*HSF1 binding, however, was poorly predictive of fly line-biased expression (between *hsal*HSF1 flies and others; Fig 5E, right, Fig S5E, left top) despite the fact that these flies showed survival differences from controls in some heat shock and aging experiments (Fig 4, Fig S4B). This may be related, in part, to the low number of DEGs biased to *hsal*HSF1 flies. We perfomed the same comparison of ChIP-seq binding with RNA-seq using the *hsal*HSF2 line relative to nRFP control flies, which yielded highly similar results (Fig S5F). Finally, we compared DEGs in *hsal*HSF2 flies also bound by *hsal*HSF2 in our fly ChIP-seq to *H. saltator* RNA-seq. Here, for both brain and fat body, ant orthologs of fly genes that were more highly expressed in *hsal*HSF2 flies showed gamergate bias among old ants (Fig S5G, top), and these genes also exhibited an age-related decrease in expression for workers but not gamergates (Fig S5G, bottom).

Taken together, we find that *hsal*HSF2 binds to and upregulates chaperone genes in control non-stress conditions in both *H. saltator* and in *D. melanogaster*; these genes contribute to increased heat stress resilience as well as to extended lifespan of both *H. saltator* gamergates and *hsal*HSF2 flies (Fig 6).

**Figure 6.**
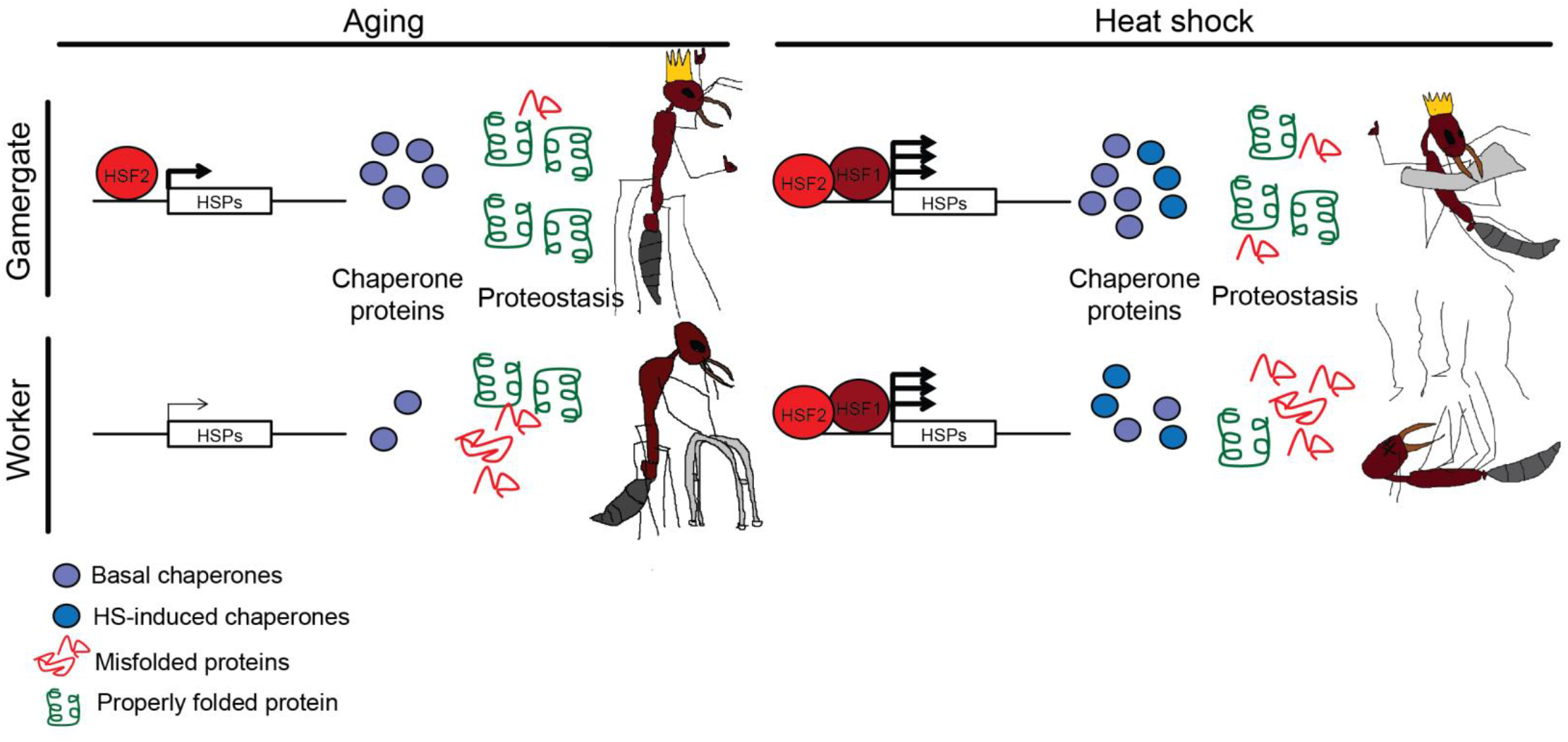
Model of *hsal*HSF2-mediated proteostasis in gamergates. Left: under normal conditions, HSF2 preferentially binds and upregulates chaperone genes in gmaergates, leading to eleveated (or maintained, age-invariant) levels of chaperone proteins, maintaining proteostasis with increased age, while workers experience the commonly-observed loss of proteostais with age. Right: upon heat stress, higher basal levels of chaperone proteins in gamergates compliment the caste-invariant heat shock response, leading to better management of heat-induced protein misfolding and increased survival of gamergates during heat stress.

## Discussion

The amazing differential lifespans occurring in the reproductive caste compared to worker castes in ant societies provides an unparalleled experimental model to uncover mechanisms underlying aging in complex organisms. The heat shock response pathway has long been investigated for transcriptional control of proteomic surveillance during both heat stress and aging stress. Our results in this study show that HSR pathway genes generally upregulated upon heat stress (in both castes) are strongly gamergate-biased in expression in basal non-stress conditions, and also increase in expression with gamergate aging. A mechanistic finding of our work that may explain the differential gene expression is that *H. saltator* encodes a gene with apparent homology to mammalian HSF2 (as do all Hymenoptera examined; Fig S2C), and that HSF2 specifically increases in expression as gamergates age, but decreases or remains lowly expressed in workers (Fig 2A). Genome-wide profiling of HSF2 and HSF1 show distinct binding patterns (Fig 2, S2G), and, further, HSF2 is bound to DNA even under basal conditions exclusively in gamergates (Figs 2D, S2E, S2G), corresponding to gamergate-biased gene expression of multiple core chaperone proteins crucial to proteostasis (Figs 1F, S2A and S2B).

A second illuminating finding of our study is that transgenic expression of *hsal*HSF2 in fly leads to improved survival under multiple heat stress regimes, and to extension of fly lifespan without heat stress; this resilience was greater than observed with similar expression of *hsal*HSF1 or *dmel*HSF (Fig 4). Profiling of the fly transcriptomes and *hsal*HSF2 binding in fly reveals conserved, distinguishing features common between transgenic flies and *H. saltator* gamergates (Figs S4C, S5, 5C, 5D), underscoring the importance of *hsal*HSF2 in accomplishing gene regulation leading to lifespan extension in gamergates via maintenance of proteome integrity. These findings have far-reaching implications due to the widely conserved link between the negative consequences of aging and breakdown of proteostasis.

The considerable lifespan disparity between reproductive and worker social insect castes provide outstanding examples demonstrating natural selection of epigenetic lifespan regulation in a natural system. *H. saltator* in particular represents an exceptional model to study this phenomenon, because any worker in the colony can transition to reproductive status (called gamergate) accompanied by a 6-fold increase in lifespan (Fig 1B). In this study we discovered that, along with lifespan extension, gamergates are buffered against heat stress-induced mortality, while workers suffer the common decrease in heat stress survival with age (Fig1A; ^3^). Thus, the gain in heat stress survival is a specific trait of the *H. saltator* gamergate state, as is lifespan of >3 years, compared to greater susceptibility of the workers to heat stress mortality and lower lifespan of ∼6-8 months. Our study thus provides insight to the underpinnings of social insect lifespan plasticity, revealing that: gamergates exhibit a strong, global transcriptomic signature of upregulating HSR genes (chaperones, stress resposnse genes) under basal conditions and with increased age; 2) *H. saltator* (and other Hymenoptera) encode an HSF2-like gene, in addition to the common HSF1 gene, and HSF2 is considerably elevated in gamergates with increased age (but declines in workers with age), 3) HSF2 shows a striking caste bias in binding under basal (non-heat stress) conditions, and this binding is predictive of stress response differential gene expression; 4) transgenic expression of *hsal*HSF2 in flies provides resiliency to heat-stress mortality, extends lifespan, and upregulates core chaperone genes.

Studies in model systems strongly link proteome “health” with longevity, because protein misfolding increases with age in many models^16,19,20^, coincident with decreasing efficiacy of the HSR^20,27^. In the short-lived worker ants, we observe considerably lower survival following heat stress than in the long-lived gamergates, and this pattern becomes more pronounced in older ants (Fig 1A), suggesting gamergates effectively manage negative consequences of protein misfolding compared to workers. This could be due to either a more rapid and elevated HSR upon heat insult, or to an overall ‘healthier’ proteome in gamergates which buffers against rapid heat-induced proteomic stress. Consistent with the latter hypothesis, we observed expression of multiple chaperone proteins that were naturally biased to gamergates in RNA-seq data taken from two tissues of young and old ants (Figs 1F, S2A and B), and generally, genes differentially regulated by heat stress in *H. saltator* correlated with genes showing gamergate-biased expression during aging in natural samples (Fig 1C, 1D, S1C and S1D). Overall these findings indicate that gamergates utilize genes typically differentially expressed upon heat stress to “repurpose” them during aging, potentially underlying the inducible extension of lifespan in gamergates. This also explains our findings that caste-biased gene expression response upon heat stress shows fewer genes preferentially up-or downregulated by gamergates—specifically that gamergates show basal elevated expression of these genes in control non-heat stress conditions, and indeed many of these genes reach similarly high level expression upon heat stress, matching levels in workers (Fig 3E, S2A and S2B).

In mammals, HSF2 is distinct from HSF1 in its activity and binding specificity ^11^. While *H. saltator*’s “HSF2” is distinct from mammalian HSF2 in lacking several C-terminal domains present in both human HSF1 and HSF2, *hsal*HSF2 shows closer sequence homology with human HSF2 compared to HSF1 or HSF4 (Fig 2B). Furthermore, our finding that *hsal*HSF2 binds without heat shock only in the gamergate genome is consistent from findings in mammals related to HSF2 binding upon upregulation ^13^. Currently the mechanistic basis of HSF2 binding and gene activation is unclear. In mammals, HSF2 forms heterotrimers with HSF1, while we detected only limited HSF1 binding without heat stress at HSF2 binding sites (and primarily in brain); we speculate that heterotrimerization may occur upon heat stress and HSF1 disassociation from Hsp90 to bind to HSF2 (Fig 2B). However, we also note that conservation of the oligimerization domain in *hsal*HSF2 is limited (Fig 2B). Further, *H. saltator* HSF1 binding is less predictive of gene upregulation upon heat stress compared to HSF2, in spite of the fact that the majority of genes bound by HSF2 are also bound by HSF1; this weaker association may be due to HSF1 binding many genes that are not upregulated during heat stress, rather than lack of targeting to heat stress upregulated genes. Hence, to summarize, in gamergates, HSF2 is selective in genome binding in non-heat stress conditions to important heat-stress chaperone genes, and how HSF2 accomplishes this remains to be determined.

Supporting a specific role of *H. saltator* HSF2 (hereafter “*hsal*HSF1/2”) in mediating enhanced response to heat stress and general aging in gamergates, ectopic expression of *hsal*HSF2 in *D. melanogaster* led to notable increases in fly survival following multiple heat shock regimes (Fig 3), including aging under mild heat (30C), and aging in the absence of heat (Fig 4E). Ectopic expression of *hsal*HSF1 and *dm*HSF provided increased survival but to a lesser extent, suggesting that overexpression of HSFs in general increases robustness to heat stress and general life-long proteomic insult, as previously seen in *C. elegans*^28,29^. As a mechanism for the improved resilience imparted by *hsal*HSF2, transcriptome sequencing of the ectopic fly lines revealed a strong distinction between *hsal*HSF2 flies and the other HSF ectopic flies (and nRFP controls) for both non-heat stressed and heat stressed flies (Fig 4). Furthermore, multiple genes elevated in basal gamergates and bound by *hsal*HSF2 in *H. saltator* showed a remarkably similar pattern of upregulation in control *hsal*HSF2 ectopic flies; indeed, among the 222 genes biased to *hsal*HSF2 flies *without* heat stress, were multiple enriched functional terms related to protein folding and chaperone activity (Fig 5A). Further, gene terms showing a similar pattern of *hsal*HSF2 fly bias *with* heat stress were enriched for DNA repair and gene expression regulation, suggesting that upon heat stress *hsal*HSF2 also causes distinct changes in gene expression which may contribute to enhanced surivival of gamergates upon heat stress.

Genome-wide profiling of *hsal*HSF1 and *hsal*HSF2 in ectopic fly lines revealed a similar contrast as in gamergates: *hsal*HSF1 is more broadly bound, while *hsal*HSF2 binds fewer genes but is, amazingly, more predictive of heat stress upregulation than *hsal*HSF1. *hsal*HSF2 was also more predictive of *hsal*HSF2 fly-biased expression, both with and without heat stress (Fig 5E and S5E), whereas, like gamergates, *hsal*HSF1 flies showed very few genes showing *hsal*HSF1-specific expression with or without heat stress (Fig 5A). How, precisely, this is achieved at the molecular level given the absence of a TAD in *hsal*HSF2 is an important question; however we propose that our consistent results in fly validate our findings in *H. saltator*, showing that *hsal*HSF2 induces chaperone gene expression, and does so in a stress- and age-protective manner. Further, why only a subset of genes bound by *hsal*HSF2 in non-heat shock conditions (in either species) show up-regulation remains to be answered, however it is very likely other mechanisms are at play that mediate the preferential up-regulation of only some genes bound by hsHSF2.

*hsal*HSF2 is unlikely to be the only factor mediating the extreme lifespan extension seen in gamergates, and may be specialized to mitigate proteomic stress associated with long lifespan in gamergates compared to workers. Indeed, while in most ants the queens are much longer lived (relative to the workers in the same species) compared to *H. saltator* gamergates^30^, queens in these other ants are typically developmentally entrenched, showing very strong, permanent changes to physiology. In *H. saltator*, in contrast, any worker can transition to gamergate during adulthood, thus there is no developmentally deep-seated differentiation. Instead, in transition to gamergate, normally short-lived workers extend lifespan up to six-fold, relying on post-developmental mechanisms. These may include conditionally-induced genes to mediate the molecular component of extended lifespan, such as upregulation of HSF to increase chaperone levels facultatively to combat proteome insult that will be pronounced in gamergates during long lifespan. However, this is only one negative consequence of aging, and hence other mechanisms will be invoked to manage other age-related degenerative factors such as transcription and epigenome regulatory instability, oxidative damage, and genome integrity. In summary, using the unique ant model system, we have identified a new major component of differential aging, and given the deep conservation in proteostasis relating to aging, these findings may have broad implications in the field of aging.

## Methods

### Harpegnathos saltator ants

All *H. saltator* used in this study originated as described in ^31^. The genetic background can be considered mixed and homogeneous, however, because ants were housed in separate colonies, colony-of-origin effects were taken into account with appropriate experimental design (treatment and control from the same colony) and statistics.

*H. saltator* colonies were housed in plastic boxes with a plaster nest chamber in a temperature (25°C) and humidity (50%) controlled ant facility on a 12 hr light/dark cycle. Ants were fed three times per week with live crickets and plaster was wet with water to prevent desiccation of the ant brood.

### Drosophila melanogaster

All flies were raised at 25°C and 50% humidity on a 12 hr light/dark cycle using standard Bloomington Drosophila Medium (Nutri-Fly). The PPL-Gal4 driver (58768) and elav-Gal4 driver (8760) lines were purchased from the Bloomington Drosophila stock center. The UAS-hsHSF1, UAS-hsHSF2, and UAS-dmHsf transgenic lines were generated by cloning the *H. saltator* or *D. melanogaster* cDNA for the respective genes into pBID-UASC^32^, followed by PhiC31 integrase-mediated transgenesis into the attP40 landing site on chromosome 2, performed by BestGene. The UAS-nRFP genetic line was generated as above, but using a donor plasmid provided by the Bonasio lab for the nRFP insert.

### RNA extraction and library preparation

*H. saltator* central brains or fat bodies were dissected from single individuals, snap-frozen and then homogenized in TriPure (Sigma). RNA was purified with typical yields of approximately to 0.6 µg total RNA per central brain or 0.9ug total RNA per fat body.

For RNA-seq, polyadenylated RNA was purified from total RNA using the NEBNext® Poly(A) mRNA Magnetic Isolation Module (NEB E7490) with on-bead fragmentation as described ^33^. cDNA libraries were prepared the same day using the NEBNext® Ultra™ II Directional RNA Library Prep Kit for Illumina® (NEB E7760). All samples were amplified using 8 cycles of PCR.

### ChIP-sequencing

For ChIP-sequencing, brains and fat bodies were dissected from seven *H. saltator* ants per replicate on ice, followed by addition of 300 µL of homogenization buffer (60 mM KCl; 15 mM NaCl; 50 mM HEPES, pH 7.5; 0.1 % Triton X-100) with 1 % formaldehyde and incubated for 5 min at 25°C with rotation. Formaldehyde was quenched using 250 mM glycine, and brains were gently passed 10–20 times through a 30G insulin syringe. The homogenate was pelleted by centrifugation for 10 min at 1,000 g at 4°C, followed by washing in homogenization buffer without formaldehyde, and re-pelleting. The homogenate was then resuspended in 130 µL lysis buffer (50 mM HEPES-KOH, pH 7.5; 140 mM NaCl; 1 mM EDTA; 0.5% Triton X-100; 0.1% Na-deoxycholate; 0.5% N-lauroylsarcosine), and sonicated with a Covaris S220 sonicator in a 130 µL micro tube for 15 min (peak incident power: 105; duty factor: 2 %; cycles/burst: 200). The lysate was cleared of insoluble material (10min at 18,000g at 4°C), the soluble fraction was diluted to 450 µL per IP (2 IPs), equalized according to DNA content (as measured with a qubit fluorometer), and 25 µL were saved as an input sonication control. A 1:1 mixture of washed and antibody-conjugated protein A and G Dynabeads (2 µg antibody per IP) were added to lysates and incubated overnight at 4°C with rotation in a final volume of 250 µL in 0.5mL tubes. The following day, antibody-bead complexes were washed twice in low-salt wash buffer (0.1 % Na-deoxycholate; 0.1 % SDS; 1 % Triton X-100; 10 mM Tris-HCl pH 8.0_RT_; 1 mM EDTA; 140 mM NaCl), once in high-salt wash buffer (0.1 % Na-deoxycholate; 0.1 % SDS; 1 % Triton X-100; 10 mM Tris-HCl pH 8.0_RT_; 1 mM EDTA; 360 mM NaCl), twice in LiCl wash buffer (0.5 % Na-deoxycholate; 0.5 % NP40; 10 mM Tris-HCl pH 8.0_RT_; 1 mM EDTA; 250 mM LiCl), and once in TE, followed by two elutions into 75 µL of elution buffer (50 mM Tris-HCl pH 8.0; 10 mM EDTA; 1 % SDS) at 65°C for 45 min with shaking (1,100 RPM). DNA was purified via phenol:chloroform:isoamyl alcohol (25:24:1) followed by ethanol precipitation. Pelleted DNA was resuspended in 25 µL TE.

Libraries for sequencing were prepared using the NEBNext Ultra II DNA Library Prep Kit for Illumina (NEB E7645), as described by the manufacturer but using half volumes of all reagents and starting material. For PCR amplification of new ChIP-seq targets, the optimal number of cycles was determined using a qPCR side-reaction using 10% of adapter-ligated, size-selected DNA. These cycle numbers were used for all subsequent replicates of the same ChIP. 13 cycles of PCR were used for Kr-h1, Met, and EcR libraries; 6 cycles were used for input controls.

For spike-in samples (2^nd^ and 3^rd^ replicates), lysate from 5 *D. melanogaster* whole flies expressing *H. saltator* HSF1 or HSF2 (10 flies total) was prepared in the same way as above (but in triple the volume), then added to 5% of *H. saltator* lysate (as determined by qubit high sensitivity DNA reagents).

### Production of custom antibodies

Antibodies against *Harpegnathos* HSF1 (residues 240-500, accession XP_011144893.1) and HSF2 (residues 105-240; XP_011141323.1) were raised against recombinant GST-tagged protein fragments. Rabbit immunization was performed by Cocalico biologics. MBP-tagged versions of the same antigens were used for affinity purification from the resulting antisera. The antibodies were validated by IP western blot (**Fig. S2D**).

### Fly crossing and rearing

All overexpression crosses (Gal4 driver > UAS-*hsal*HSF1/*hsal*HSF2/*dm*Hsf/nRFP) for heat shock survival, RNA-seq and ChIP-seq were reared at 25°C and 50% humidity on a 12-hour light/dark cycle using standard Bloomington Drosophila Medium (Nutri-Fly). Day0-1 F1 female flies were separated and reared in new vials. For aging, 10 female flies per vial were kept at either 25°C or 30°C with 12hr light/dark cycle, in vials with 5mL of the standard Bloomington *Drosophila* Medium (Nutri-Fly). Flies were flipped three times per week, and mortality was recorded. For the long-term heat shock survival assay, 10 female flies (age 2-5 days) per vial were incubated at 35.5°C for 24h, removing vials every 4h for 24h. Heat shocked flies were kept at 25°C for 24h before assessing survival. For shorter heat shock survival assays, 10 female flies (age 2-5 days) per vial were incubated at the given temperature (Figure 4), removed after heat shock followed by mortality assessment the following day.

### Fly Climbing Assay

Fly climbing assay was performed mainly following previous method^34^. 10 female flies of each genotype were transferred to a new vial on the given day (d19 for elav-Gal4 crossed flies, d21 for PPL-Gal4 crossed flies). 3 taps were given to knock all flies in the chamber to the bottom. After 10s of climbing, % of flies not passing the drawn line was recorded.

### Fly RNA-seq and ChIP-seq samples

For RNA-seq and ChIP-seq, female F1 offspring of UAS overexpression lines crossed to PPL-Gal4 flies were separated at d0-1. At d7, heat shock flies were incubated at 42°C for 30min, followed by 15min of recovery then placement on ice, along with control flies. For RNA-seq, a single whole female fly was used for each replicate, homogenized in TriPure (Sigma), followed by RNA-precipitation and poly-A selection for mRNA-seq.

For ChIP-seq samples, 3 whole female flies were homogenized in 500µL of homogenization buffer (60 mM KCl; 15 mM NaCl; 50 mM HEPES, pH 7.5; 0.1 % Triton X-100) with 1 % formaldehyde and incubated for 5 min at 25°C with rotation. Formaldehyde was quenched using 250 mM glycine. The homogenate was pelleted by centrifugation for 10 min at 1,000 g at 4°C, followed by washing in homogenization buffer without formaldehyde, and re-pelleting. The homogenate was then resuspended in 1mL lysis buffer (50 mM HEPES-KOH, pH 7.5; 140 mM NaCl; 1 mM EDTA; 0.5% Triton X-100; 0.1% Na-deoxycholate; 0.5% N-lauroylsarcosine), rotated for 15minutes at 4C, and sonicated with a Covaris S220 sonicator. The lysate was cleared of insoluble material (10min at 18,000g at 4°C), 600µL of the soluble fraction was taken per IP (2 IPs), equalized according to DNA content (as measured with a qubit fluorometer), and 25 µL were saved as an input sonication control. A 1:1 mixture of washed and antibody-conjugated protein A and G Dynabeads (2 µg antibody per IP) were added to lysates and incubated overnight at 4°C with rotation in a final volume of 250 µL in 0.5mL tubes. The following day, antibody-bead complexes were washed twice in low-salt wash buffer (0.1 % Na-deoxycholate; 0.1 % SDS; 1 % Triton X-100; 10 mM Tris-HCl pH 8.0_RT_; 1 mM EDTA; 140 mM NaCl), once in high-salt wash buffer (0.1 % Na-deoxycholate; 0.1 % SDS; 1 % Triton X-100; 10 mM Tris-HCl pH 8.0_RT_; 1 mM EDTA; 360 mM NaCl), twice in LiCl wash buffer (0.5 % Na-deoxycholate; 0.5 % NP40; 10 mM Tris-HCl pH 8.0_RT_; 1 mM EDTA; 250 mM LiCl), and once in TE, followed by two elutions into 75 µL of elution buffer (50 mM Tris-HCl pH 8.0; 10 mM EDTA; 1 % SDS) at 65°C for 45 min with shaking (1,100 RPM). DNA was purified via phenol:chloroform:isoamyl alcohol (25:24:1) followed by ethanol precipitation. Pelleted DNA was resuspended in 25 µL TE. ChIP-seq libraries were prepared as for those from ant tissues (above).

### Statistical analyses

#### Assignment of gene orthology and functional terms

Genes (NCBI *Harpegnathos saltator* NCBI annotation release 102; assembly v 8.5) were assigned orthology using the reciprocal best BLAST hit method ^35^ to both *D. melanogaster* (r6.16) and *H. sapiens* (GRCh38) protein coding genes. These corresponding relationships are provided in^36^.

GO terms were assigned to genes using the blast2go tool ^37^ using the nr database, as well as InterPro domain predictions. GO enrichment tests were performed with the R package topGO (Alexa and Rahnenfuhrer, 2020 ^38^, utilizing Fisher’s elimination method, and resulting significant terms were entered into REVIGO ^39^ for collapsing of redundant terms.

#### RNA-seq analysis

Reads were demultiplexed using bcl2fastq2 (Illumina) with the options “--mask-short-adapter-reads 20 --minimum-trimmed-read-length 20 --no-lane-splitting --barcode-mismatches 0”. Reads were aligned to the *H. saltator* v8.5 assembly ^40^ using STAR ^41^. STAR alignments were performed in two passes, with the first using the options “--outFilterType BySJout --outFilterMultimapNmax 20 --alignSJoverhangMin 7 -- alignSJDBoverhangMin 1 --outFilterMismatchNmax 999 --outFilterMismatchNoverLmax 0.07 -- alignIntronMin 20 --alignIntronMax 100000 --alignMatesGapMax 250000”, and the second using the options “- -outFilterType BySJout --outFilterMultimapNmax 20 --alignSJoverhangMin 8 --alignSJDBoverhangMin 1 -- outFilterMismatchNmax 999 --outFilterMismatchNoverLmax 0.04 --alignIntronMin 20 --alignIntronMax 500000 --alignMatesGapMax 500000 --sjdbFileChrStartEnd [SJ_files]” where “[SJ_files]” corresponds to the splice junctions produced from all first pass runs.

Gene-level read counts were produced using featureCounts ^42^ with the options “-O -M --fraction -s 2 – p”, against a custom adapted NCBI *Harpegnathos* annotation (see following paragraph). Resulting counts were rounded to integers prior to importing into R for DESeq2 analysis in order to account for fractional count values associated with fractional counting of multi-mapping reads.

Because we observed a large number of genes which possessed truncated 3’UTRs in the *H. saltator* NCBI annotation relative to aligned brain RNA-seq data, we sought to globally amend annotations to account for this. In order to create this adapted NCBI annotation, we first aligned all reads as above, but in the second round of alignment included the option “--outSAMattributes NH HI AS nM XS”. We then ran isoSCM v2.0.12 ^43^ assemble with the options “-merge_radius 150 -s reverse_forward” on all alignments. Following this we extracted only 3’ exons from the resulting draft annotations, which were then overlapped with the current NCBI annotation release. We ten took the longest predicted 3’ exon/UTR that overlapped with the current NCBI version’s 3’ exon, and extended this exon according to the isoSCM 3’ exon. Any predicted 3’ exon extension that overlapped an exon of another gene on the same strand was trimmed to 500 bp upstream of the other gene’s offending exon.

Differential gene expression tests were performed with DESeq2 ^44^. For all pairwise comparisons the Wald negative binomial test (test = “Wald”) was used for determining differentially expressed genes. For caste-specific changes with condition (e.g. caste-specific aging changes, or caste-specific heat shock response changes in gene expression) the specific caste X condition was compared to all other samples in a given comparison after blocking for colony background and general caste effects. When testing for general heat shock DEGs samples from both castes were utilized, comparing HS to control samples after blocking for caste. Unless otherwise stated, an adjusted *P*-value cutoff of 0.1 was used to define differentially expressed genes.

For the PCA analysis in Figure 5B, in order to identify genes differing across samples in all contexts we performed a likelihood ratio test in DESeq2 comparing the full model ‘∼line+condition+line:condition’ to the reduced model ‘∼1’. Genes with an adjusted p-value < 0.01 were used to perform a PCA on variance stabilizing transformed expression values.

#### ChIP-seq analysis

Reads were demultiplexed using bcl2fastq2 (Illumina) with the options “--mask-short-adapter-reads 20 - -minimum-trimmed-read-length 20 --no-lane-splitting --barcode-mismatches 0”. Reads were trimmed using Trimmomatic ^45^ with the options “ILLUMINACLIP:[adapter.fa]:2:30:10 LEADING:5 TRAILING:5 SLIDINGWINDOW:4:15 MINLEN:15”, and aligned to the *Harpegnathos* v8.5 assembly ^40^ using bowtie2 v2.2.6 ^46^ with the option “--sensitive-local”. Alignments with a mapping quality below 5 and duplicated reads were removed using samtools ^47^, as were reads overlapping a custom set of blacklist regions, produced as in ^48^, using input sonication control samples from this study, as well as ^36^. Peaks were called using macs2 v2.1.1.20160309 ^49^ with the options “--call-summits --nomodel -B”.

To identify general HSF1 and HSF2 peaks, reads from all heat shocked sample replicates, regardless of caste, were used. For HSF2 reads from gamergate control samples were also included. Only peaks with a fold-enrichment over input > 3 were considered, in order to produce a list of high-confidence peaks to associate to genes. For peak classification, promoters were defined as the region spanning 2 kb upstream and 0.5 kb downstream the transcription start site of a gene. Genes bound by HSFs were defined as those genes containing a “strong” (fold-enrichment over input > 3) ChIP-seq peak within the promoter.

Differential HSF ChIP peaks (Figs 2, 3, S5, and S3) were called using DiffBind ^50^ utilizing the option “summits=250” in the dba.count() function, and “bFullLibrarySize=FALSE, bSubControl=TRUE, bTagwise=FALSE” for dba.analyze() and all peaks called by macs2 were used regardless of fold enrichment. For comparisons between where one condition largely lacked any peaks (e.g. heat shock vs control samples, as well as HSF2 caste-specific binding without heat shock) the option “bFullLibrarySize=TRUE” was used, in order to avoid normalization artifacts arising from scaling of the background read counts in the unbound condition.

For DiffBind testing the DESeq2 algorithm (Wald negative binomial test) with blocking was used, and ChIP replicate was used as the blocking factor while testing for caste differences. When comparing general heat shock vs control peaks caste was used as a blocking factor. An adjusted *P*-value of 0.1 was used as threshold to define differential binding.

To generate genome browser tracks and heatmaps, RPM (reads per million) were calculated for each biological replicate and then averaged.

For determination of general peaks and HSF-bound genes (Fig S2E, S4A), peaks were called in individual replicates, and a peak was retained if independently called in >2 replicates.

#### Statistics

Sample size and statistical tests are indicated in the figure legends. Unless otherwise noted all statistical tests were two-sided. Boxplots were drawn using default parameters in R (center line, median; box limits, upper and lower quartiles; whiskers, 1.5x interquartile range). For testing significance of gene overlaps, we used the GeneOverlap R package ^51^ and the R package OrderedList^52^. For comparisons given in boxplots, non-parametric tests were used: Mann-Whitney U tests were used for all one-way comparisons, and Kruskal-Wallis test followed by Dunn’s correction for comparisons between 3 or more groups.

## Acknowledgements

We thank members of the Berger Lab for help in editing the manuscript. This work was supported by NIH training grant F32GM120933 (K.M.G.) and NIA R01 5R01AG055570 (S.L.B.).

## Supplementary Figures

**Figure S1:**
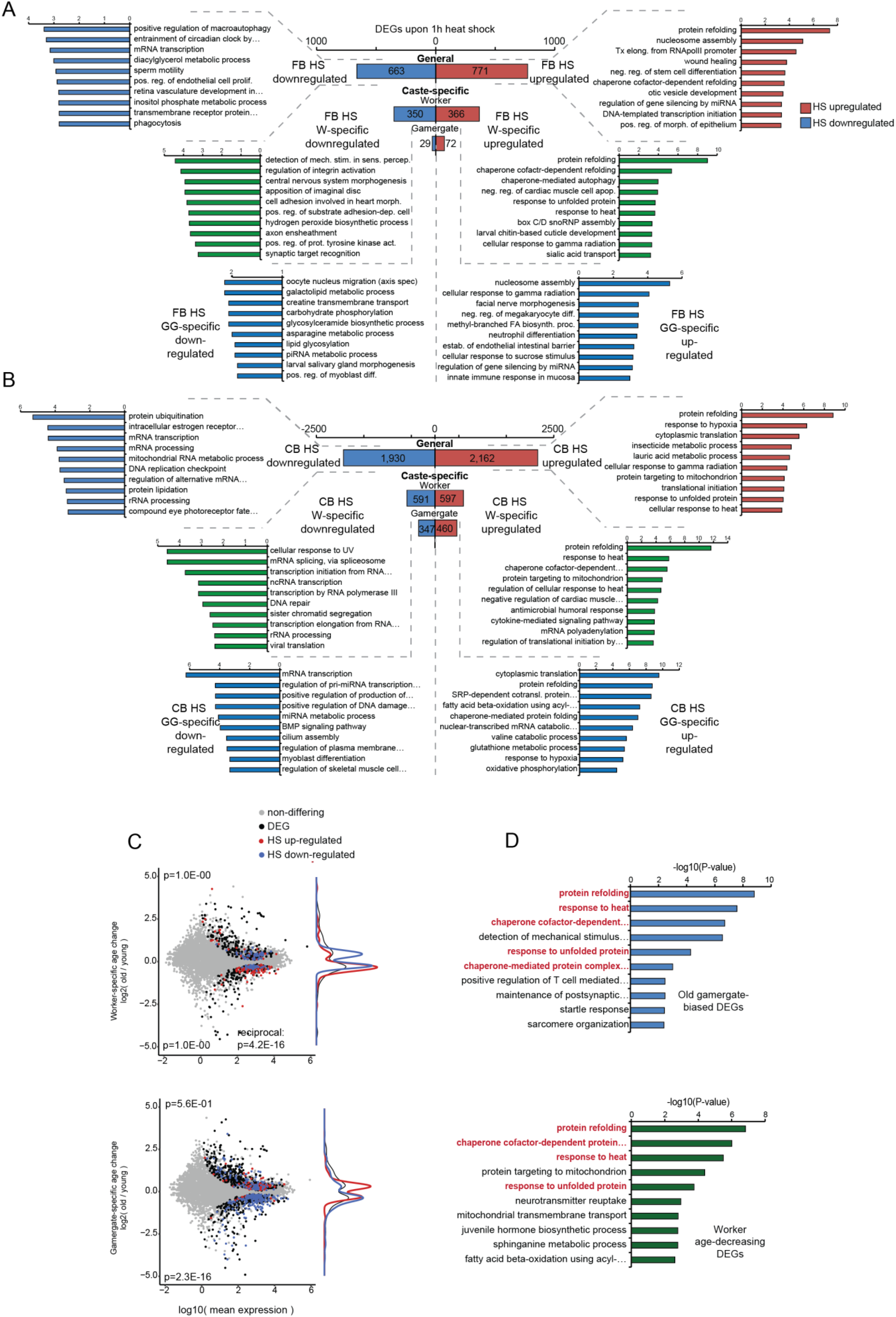
A and B) Number of genes showing differential expression upon heat shock between both castes (analyzed together; “General”), as well as those showing Worker or Gamergate preferential up-or down-regualtion (see methods), illustrating strikingly fewer genes are preferentially up-regulated in gamergates 1h following heat shock. Given beside each are bar plots showing top ten gene ontology terms enriched among each category for (A) Fat body heat shock DEGs, and B) Brain heat shock DEGs. C) MA plots of genes showing caste-specific age-related changes in expression, with 1h HS-up and HS-down regulated genes colored in red and blue respectively. P-values in upper and lower are from a fishers exact test comparing up- and down-regulated caste-biased age changing genes to 1h heat shock up- and down-regulated DEGs respectively. D) Gene ontology term (Biological Process) results for the top ten terms enriched among genes (top) biased to old gamergate brains as compared to old workers and (bottom) showing significant decrease in worker brainss relative to gamergates with age, illustrating enrichment of proteome and chaperone related terms in these categories.

**Figure S2:**
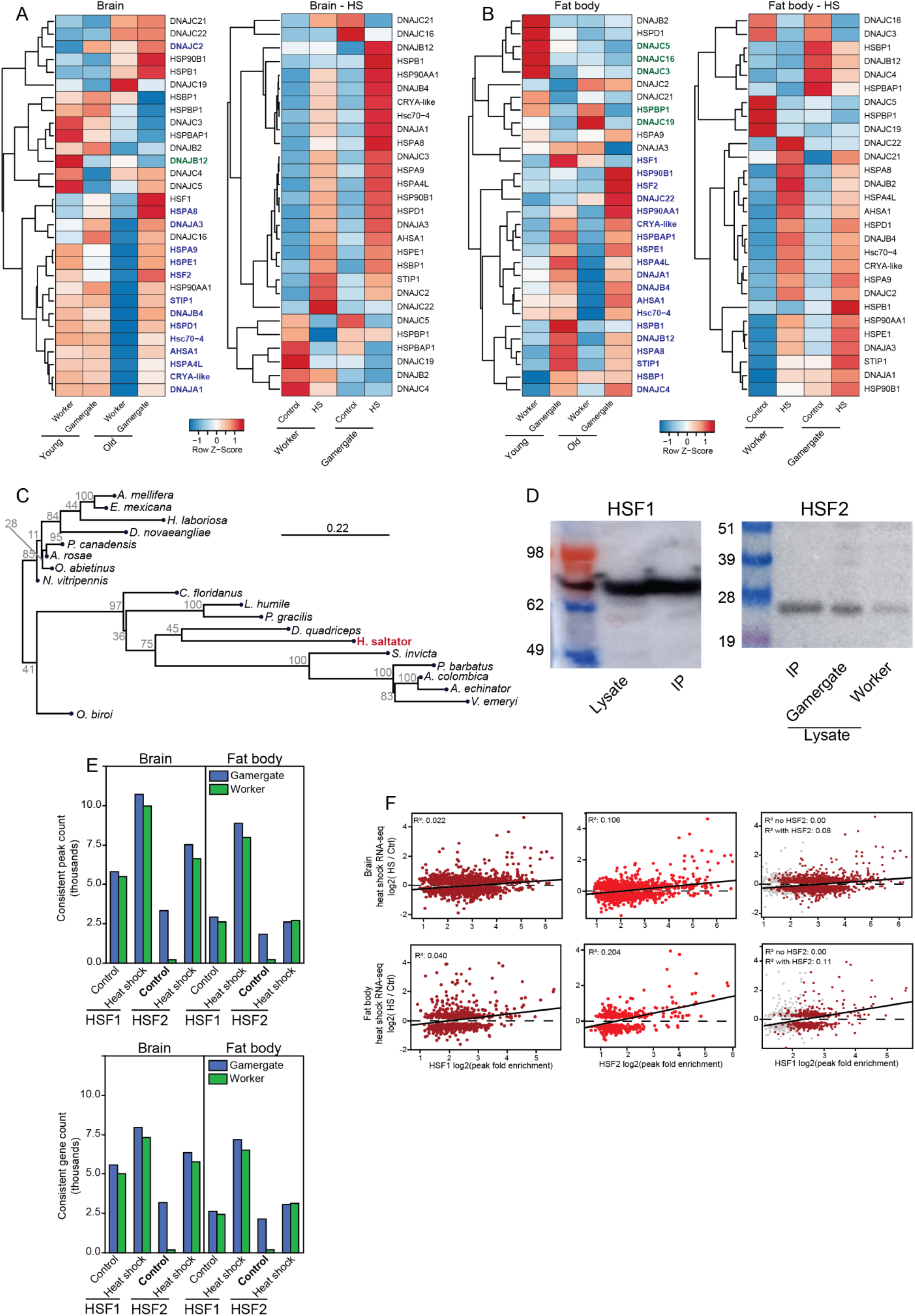
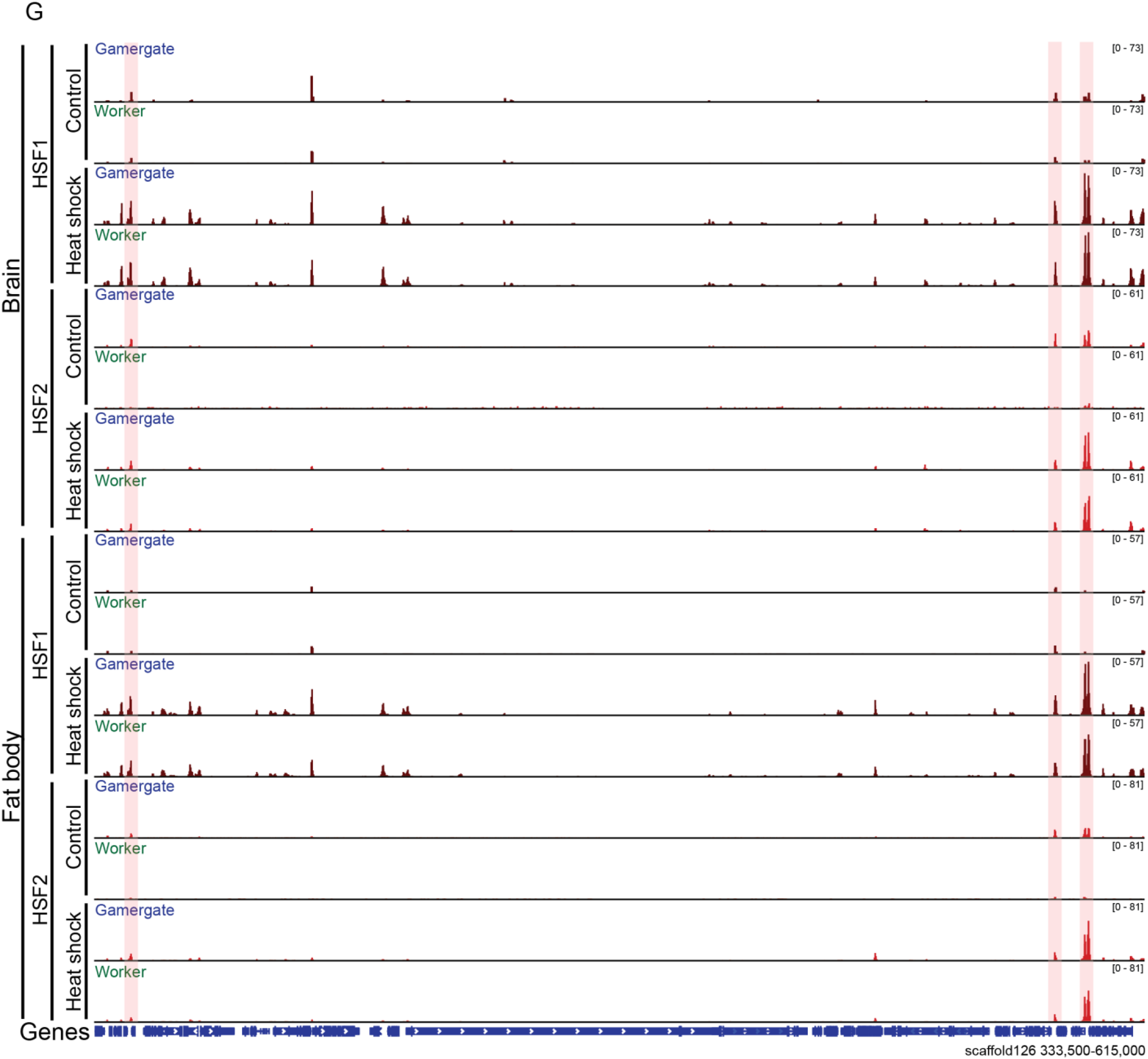
A) Heatmap of RNA-seq expression in brain for 28 genes with homology to known HSP or chaperone genes in human or fly for aging RNA-seq (left) as well as heat stress RNA-seq (right). Genes are colored if significantly biased to gamergate (blue) or worker (green) in aging brain aging RNA-seq data when comparing all gamergates to workers (blocking for age) or when comparing old gamergates to old workers. Data represents row z-scores from normalized counts. B) The same as in S2A, but for RNA-seq data taken from fat body. C) Phylogenetic tree generated from protein alignment (PRANK) of HSF2 genes from representative species within Hymenoptera (4 bees, 4 wasps, 11 ants). Branch lengths represent protein sequence divergence (number of AA changes per site). Numbers at each node represent bootstrap values generated from 10,000 permutations. D) IP-western blot validation for antibodies targeting HSF1 (left) and HSF2 (right). For HSF2 lysate from d250 worker and gamergate fat bodies (10% of 10x each, without heat shock) were run to illustrate different protein levels in each caste, then pooled for IP. E) Number of peaks (left) and number of peak-bound genes (right) for HSF1 and HSF2 in worker and gamergate, illustrating very low peak count for HSF2 in worker without heat shock (Control), while non-heat shocked gamergates show similar numbers of peaks with and without heat shock. F) Correlations between HSF peak enrichment level and general heat shock gene expression changes (both castes combined) for (left) HSF1, (middle) HSF2, and HSF1 at HSF2 peaks (right), for either brain (top row) or fat body (bottom row). Presented are genes showing significant differential gene expression comparing heat shock and control (padj < 0.1) and a peak of the given HSF in at least two replicates upon heat shocked conditions. G) ChIP-seq tracks from brain (top tracks) and fat body with and without heat shock for both castes, showing a ∼300kb window, illustrating more binding by HSF1, as well as binding of HSF2 in gamergates (red highlighted regions) under basal conditions. Values given in upper right of each track represent data range (RPM).

**Figure S3:**
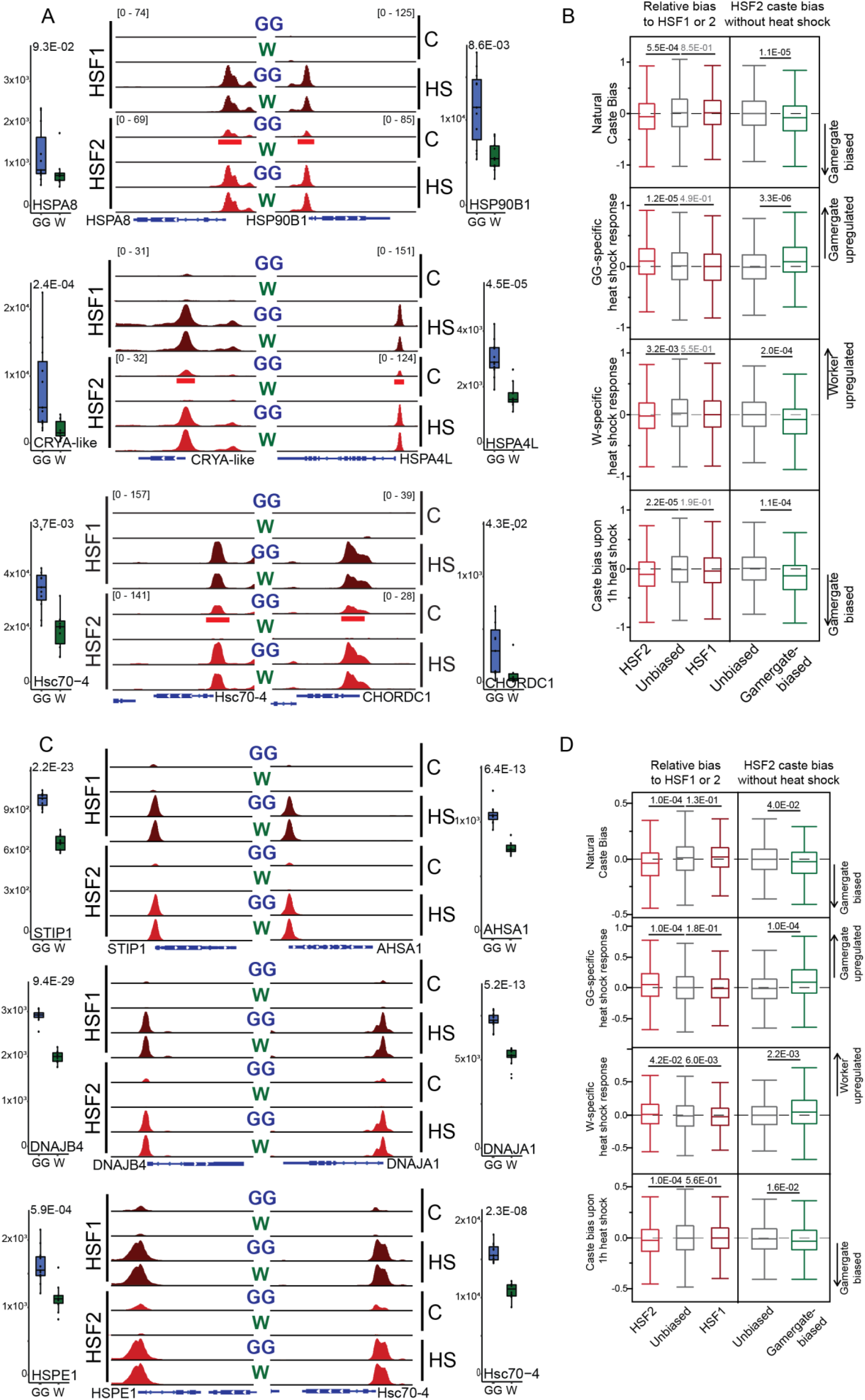
ChIP-sequencing of HSF1 and HSF2 in ants reveals HSF- and caste differing binding. A) Example genome browser tracks for 6 more genes (related to Fig 2E) shown in Fig 1F further illustrating binding by HSF2 in gamergates in the absence of heat shock (red underlined peaks). For respective HSFs, all tracks were autoscaled to one another (RPM values given in brackets). C: Control, HS: heat shock, GG: gamergate samples, W: worker samples. RNA-seq expression count plots given aside each track representing RNA-seq data from old gamergates (blue) and workers (green). B) The same analysis as in Figure 2E, but for all genes. C) Example genome browser tracks for 6 genes matching those in Figure 2F, but representing data from brain illustrating similar pattern of HSF2 binding without heat shock only in gamergates. For respective HSFs, all tracks were autoscaled to one another (RPM values given in brackets). C: Control, HS: heat shock, GG: gamergate samples, W: worker samples. RNA-seq expression count plots given aside each track representing RNA-seq data from old gamergates (blue) and workers (green). D) The same plot as for S3B above, but using data from brain, illustrating conserved patterns as seen in fat body but for brain.

**Figure S4.**
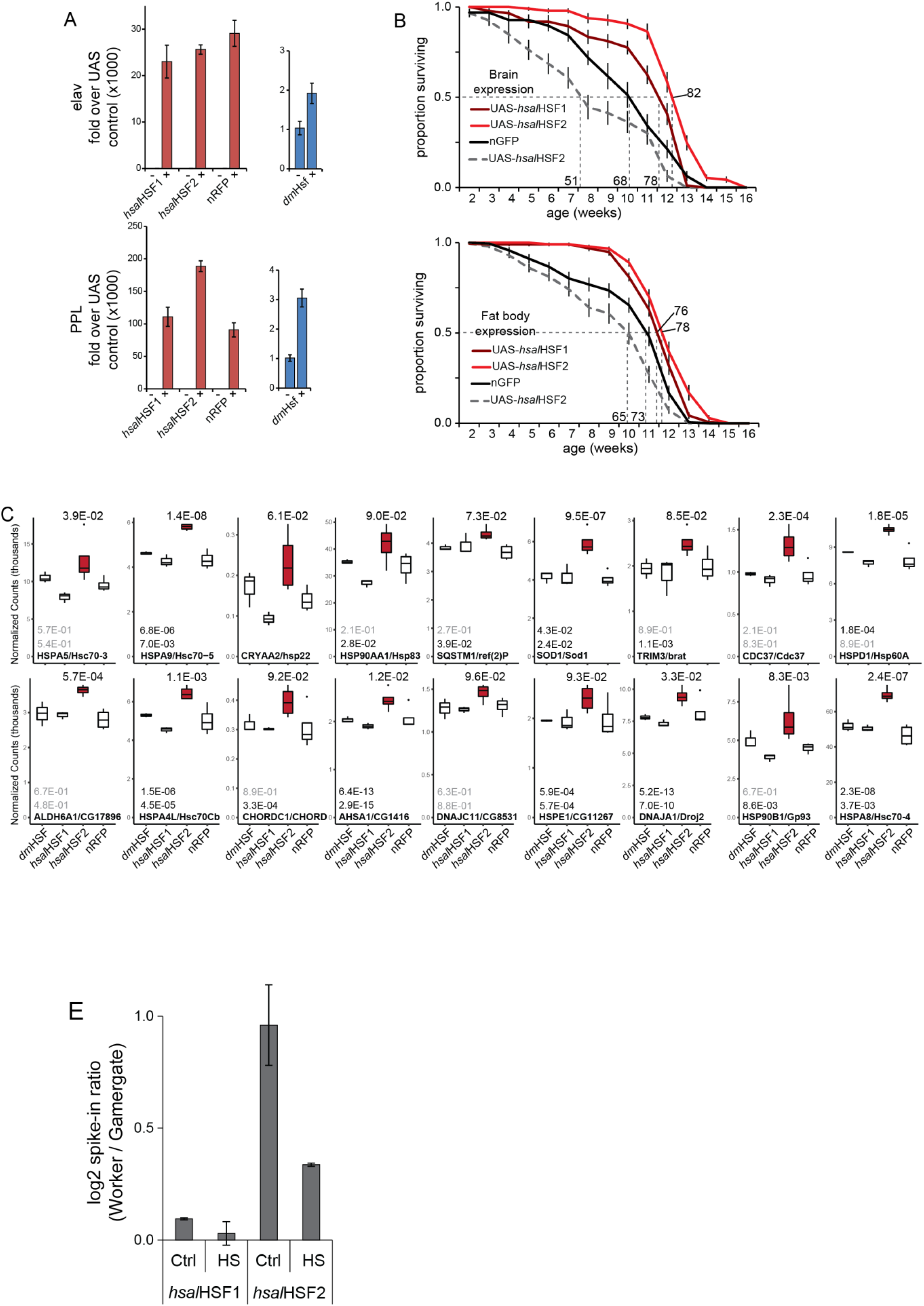
*D. melanogaster* ectopic fly results. A) Validation of fly ectopic expression lines via RT-qPCR. Shown are relative fold expression levels of flies crossed to Gal4-elav (top) or Gal4-PPL (bottom) driver lines (+ samples) as compared to Parental UAS lines (-samples) for hsHSF as well as nRFP control lines. In blue the same is given for UAS-dmHSF, plotted separately due to endogenous expression of this gene. B) Results from secondary general aging experiment (similar to Figure 4E) comparing HSF ectopic fly line survival under standard conditions. Using nGFP (iso31 genetic background) and UAS-*hsal*HSF2 (without gal4 driven expression) as controls. Numbers attached to each curve represent median fly surivival in days. C) normalized count plots showing genes with chaperone functions that are preferentially elevated in hsHSF2 ectopic flies in the absence of heat shock, illustrating similarity with that seen in *H. saltator* gamergates. Values given above gene names represent adjusted p-values taken from *H. saltator* caste comparisons between old gamergates and workers for gamergate-biased genes. top: central brain adjusted p-values; bottom: fat body adjusted p-values. If no adjusted p-values given (hsp22) this gene does not have a reciprocal best hit in *H. saltator*. E) log2-transformed ratio of worker:gamergate input-normalized spike-in proportions from ChIP-seq of hsHSF1 and hsHSF2 in *H. saltator* using a pool of lysate from hsHSF1 and hsHSF2 *D. melanogaster* ectopic expression fly lines as spike-in. Positive values indicate more spike-in present in worker ChIP samples, indicating less endogenous DNA-bound protein. Error bars represent standard deviation (n=2).

**Figure S5.**
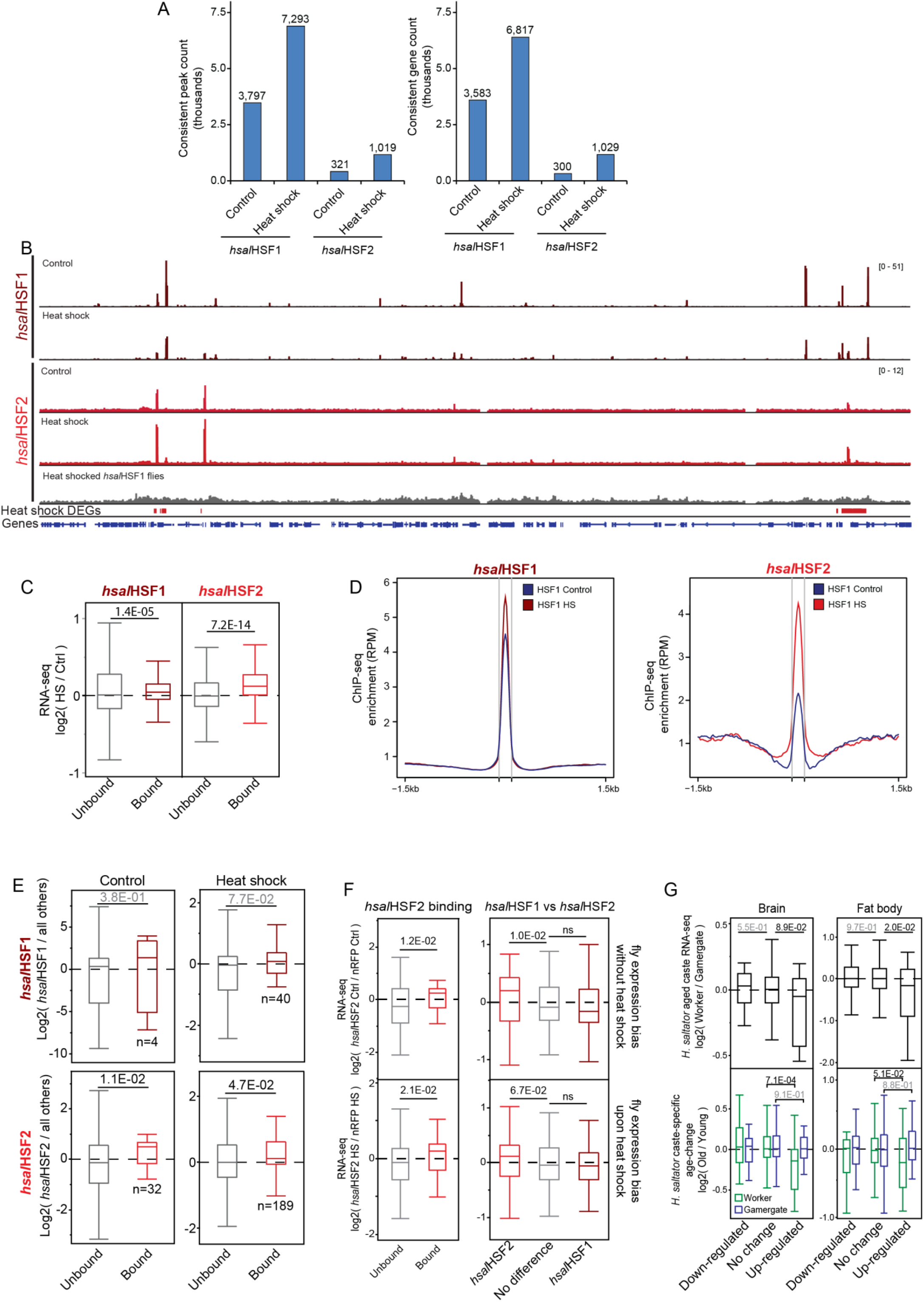
ChIP-sequencing of hsHSFs in fly lines. A) Number of peaks (left) and peak-marking genes (right) detected with each fly line X ChIP experiment in both conditions. B) Example genome browser view of a 600kb (r6.42, chr3L:7,800kb-8,400kb) region from ectopic flies expressing hsHSF1 (maroon) and hsHSF2 (red) with and without heat shock illustrating that contrasting heat shock and control samples hsHSF1 binds in a distinct manner as compared to hsHSF2. Also given (grey track) is ChIP-seq using hsHSF2 antibody in UAS-hsHSF1 (PPL-Gal4 driven) flies, illustrating lack of cross-reactivity. C) Similar to that seen in *H. saltator* (Figure 2C) hsHSF2 binding in transgenic flies is more predictive of heat shock up-regulation than hsHSF1. Presented are log2 fold changes between control and heat shocked fly RNA-seq samples for the respective hsHSF, comparing heat shock peak-marked genes to those without a peak for the respective hsHSF. P-values represent a mann-whitney U test between unbound and bound genes. D) Metaplots of average ChIP-seq signal (RPM-normalized signal) at peaks for the respective HSF without (blue) and with (red) heat shock, illustrating that hsHSF2 shows greater dynamicism between heat shocked and non-heat shocked conditions vs HSF1. E) Boxplots comparing hsHSF bound and unbound genes differentially expressed between hsHSF1 (top) and hsHSF2 (bottom) fly lines and all others in this study, both among control flies (left plots) and heat shocked flies (right plots), illustrating that genes bound by hsHSF2 are biased in expression to hsHSF2 flies, while hsHSF1 does not have this relationship. P-values from a mann-whitney U test comparing DEGs unbound by the given hsHSF to those bound by the given hsHSF. F) Related to Figure 4E, boxplots of average log2 expression ratios for DEGs between HSF2 flies and nRFP flies, illustrating that (left) HSF2 binding (as determined by peak presence in ChIP-seq of transgenic flies) is predictive of HSF2 fly transcriptional bias when only comparing to negative control ectopic flies both for samples in the absence of heat shock (top) and heat shocked samples (bottom), as is (right) higher relative enrichment of HSF2 (vs hsHSF1). P-values from a Mann-whitney U test comparing genes without HSF2 or with no difference between HSF2 and HSF1 to the respective categories. G) Boxplot showing (top) level of caste bias in *H. saltator* RNA-seq data taken from old workers and gamergates and (bottom) worker and gamergate-specific aging changes in expression for (left) brain and (right) fat body, plotted at orthologous fly genes showing either up (n=26) or downregulation (n=19) in *D. melanogaster* fly lines to *hsal*HSF2 and featuring *hsal*HSF2 DNA binding in flies without heat shock, illustrating that genes up-regulated in *hsal*HSF2 flies without heat shock show gamergate bias in *H. saltator* and decrease in expression in aging workers but not gamergates. P-values from a mann-whitney U test comparing up- and down-regulated genes in *hsal*HSF2 flies vs unchanging genes.

## Notes

### Competing Interest Statement

The authors have declared no competing interest.

